# Folate prevents the autism-related phenotype caused by developmental pyrethroid exposure in prairie voles

**DOI:** 10.1101/2024.11.25.625285

**Authors:** Nilanjana Saferin, Ibrahim Haseeb, Adam M. Taha, Sarah E. Beecroft, Sangeetha Pillai, Asha E. Neifer, Rudhasri Lakkuru, Brian P. Kistler, Charlotte N. Nawor, Isa Malik, Dena Hasan, Jonathan A. Carlson, Kareem K. Zade, Sydnee P. Dressel, Eileen M. Carney, Radha Shah, Shudhant Gautam, John Vergis, Kari L. Neifer, Zachary V. Johnson, Morgan L. Gustison, F. Scott Hall, James P. Burkett

## Abstract

Neurodevelopmental disorders (NDDs) have dramatically increased in prevalence to an alarming one in six children, and yet both causes and preventions remain elusive. Recent human epidemiology and animal studies have implicated developmental exposure to pyrethroid pesticides, one of the most common classes of pesticides in the US, as an environmental risk factor for autism and neurodevelopmental disorders. Our previous research has shown that low-dose chronic developmental pyrethroid exposure (DPE) changes folate metabolites in the adult mouse brain. We hypothesize that DPE acts directly on molecular targets in the folate metabolism pathway, and that high-dose maternal folate supplementation can prevent or reduce the biobehavioral effects of DPE. We exposed pregnant prairie vole dams chronically to vehicle or low-dose deltamethrin (3 mg/kg/3 days) with or without high-dose folate supplementation (methylfolate, 5 mg/kg/3 days). The resulting DPE offspring showed broad deficits in five behavioral domains relevant to neurodevelopmental disorders (including the social domain); increased plasma folate concentrations; and increased neural expression of SHMT1, a folate cycle enzyme. Maternal folate supplementation prevented most of the behavioral phenotypes (except for repetitive behaviors) and caused potentially compensatory changes in neural expression of FOLR1 and MTHFR, two folate-related proteins. We conclude that DPE causes neurodevelopmental disorder-relevant behavioral deficits; DPE directly alters aspects of folate metabolism; and preventative interventions targeting folate metabolism are effective in reducing, but not eliminating, the behavioral effects of DPE.

## INTRODUCTION

The prevalence of neurodevelopmental disorders (NDDs) in the US has dramatically increased in the past few decades to an alarming one in six children [1]. This crisis is particularly concerning considering that most NDDs lack effective medical treatments or reliable biomarkers and can only be diagnosed behaviorally [2]. Effectively addressing this crisis will require addressing the critical lack of knowledge regarding causes and effective prevention strategies.

Although NDDs are traditionally considered to be primarily genetic, the risk contributed by environmental factors has been more recently estimated at 38-58% [3–5]. Similarly, the dominant focus of animal research relevant to NDDs has traditionally been on models of genetic risk [6, 7]. Nonetheless, some environmental risk factors for NDDs have been identified and studied, including lifestyle factors (such as parental age) [8], internal environment (maternal immune response, prenatal hormone exposure, and others) [9–15], and developmental exposures (medications, chemical exposures, and others) [16–22]. Additional research is critically needed to evaluate the causal roles and biological mechanisms of these environmental risk factors.

One environmental factor of interest is developmental exposure to pyrethroid pesticides. Pesticides are biocidal compounds often used as household insecticides [23] and are essential for agricultural purposes and public health management of mosquitos [24–26]. Because of their supposed safety in adults, pyrethroid insecticides are among the most frequently used pesticides in the US [27]. Due to their extensive usage, pyrethroids are universally present in urban streams and runoff water [28, 29] and are detectable in blood in 70-80% of the US population [30, 31] and 99.7% of pregnant women in France [32]. Nonetheless, evidence from human research suggests a link between prenatal exposure to pyrethroids and risk for NDDs [22, 25, 30, 31, 33–38]. Mouse models of developmental pyrethroid exposure have provided evidence that this low-dose exposure may be a cause of NDD-relevant behavioral and neuromolecular phenotypes [30, 39, 40].

One study has suggested that high levels of folate supplementation (>800 µg/day) in pregnant women is correlated with a reduction in the risk for NDDs contributed by pyrethroid exposure [41]. Critically, our previous research in mice showed an effect of developmental pesticide exposure on non-specific folate metabolites in the brain [42]. These findings suggest an important and, as of yet, poorly understood link between folate metabolism and pyrethroid exposure.

We hypothesize that developmental pyrethroid exposure exerts its adverse behavioral and neuromolecular effects, in whole or in part, through direct interaction with folate metabolism. Further, we hypothesize that maternal folate supplementation, specifically using the biologically active form of folate (methylfolate, also known as 5-methyltetrahydrofolate, 5-MTHF, or levomefolic acid) to bypass disruptions in folate metabolism, reduces or prevents these adverse behavioral and neuromolecular effects.

To test these hypotheses, we exposed prairie vole dams orally to a chronic low dose (3 mg/kg or vehicle every 3 days) of the EPA reference pyrethroid deltamethrin [43] during pre-conception, pregnancy, and lactation, with or without high dose methylfolate supplementation (5 mg/kg).

Prairie voles (*Microtus ochrogaster*) are wild rodents indigenous to the Midwest that are known for unique characteristics that provide distinct advantages in research related to neurodevelopmental disorders, including their complex social behaviors; their outbred genetics representative of the wild; and their individual variability that can be linked to biological variability [44–49]. We predicted that the prairie vole’s species-specific complex behaviors, including monogamous pair bonding and empathy-based consoling behavior, would be more sensitive to developmental disruption than the relatively simplistic and fundamental social behaviors of mice. We tested the resulting offspring on a wide range of behavioral outcomes in five behavioral domains relevant to NDDs: the social, communication, cognitive, locomotor, and repetitive behavior domains [50]. We also examined the brains of adult offspring for specific changes in the folate metabolism pathway caused by developmental exposure with or without supplementation. In accordance with gold-standard recommendations from behavioral neuroscientists working on NDDs [2], we maximized rigor and reproducibility using an animal model possessing complex social behaviors; a broad NDD-relevant behavioral domain approach; multiple tests within each behavioral domain; maximization of automated coding; testing at multiple life stages; a litter-based design; and male and female subjects.

## MATERIALS AND METHODS

### Animals

All subjects used for our study were healthy male and female prairie voles (*Microtus ochrogaster*) from our wild-type breeding colony at the University of Toledo. Our prairie vole colony breeders were obtained from the Cornell Vole Core, whose stock was derived from wild-caught animals captured at field sites in Illinois. All wild-type prairie voles in our colony were within five generations of their wild ancestors. Colony breeders were outbred in monogamous pairs from unrelated lineages, and their offspring were weaned into same-sex cages housing 2-3 voles at 21 days of age. All voles were maintained on a 12-hour light/dark cycle and provided with tap water and rabbit chow (5326, LabDiet, St. Louis, MO) ad libitum. Importantly, LabDiet rabbit chow contains 7.4 ppm “Folic Acid,” which LabDiet reports is 81% in added synthetic folic acid form and 19% in dietary folates from natural ingredients [51]. The voles were housed in ventilated rat cages outfitted with paper bedding, nesting enrichment and chewing enrichment as previously described [52] (cages: Allentown, Allentown, NJ; bedding: Pure o’Cel, The Andersons, Maumee, OH; nesting: Bed r’Nest, The Andersons, Maumee, OH; nestlets: Ancare, Bellmore, NY; crawl balls: Bio-Serv, Flemington, NJ; chewing: 500 g Apple Sticks, Amazon, Seattle, WA). All procedures were approved by the University of Toledo IACUC and conducted in compliance with the Animal Welfare Act and the National Research Council’s Guide for the Care and Use of Laboratory Animals.

### Study design

Our study design (Fig. 1A) consisted of three groups: vehicle exposure (Control), developmental pyrethroid pesticide exposure (DPE), and pesticide exposure supplemented with the folate vitamer methylfolate (DPE+F). Prairie vole dams were given a chronic low-dose exposure to deltamethrin (3 mg/kg/3 days), vehicle, or deltamethrin plus high-dose methylfolate (5 mg/kg/3 days) during preconception, pregnancy, and lactation as described below. After two weeks of pre-conception exposure, the dams were separated from their same-sex cage mates and permanently pair-housed with a male sire from an unrelated lineage. The sire remained in the cage throughout pregnancy and lactation to provide species-typical biparental care to the offspring. Male and female offspring were considered the subjects of the study and were used for subsequent experiments. Experimental measures on offspring began at 1 week of age (ultrasonic vocalizations) and continued through adulthood. Offspring were weaned at 21 days of age and housed in same-sex cages of 2-3 voles from the same experimental group.

**Figure 1.**
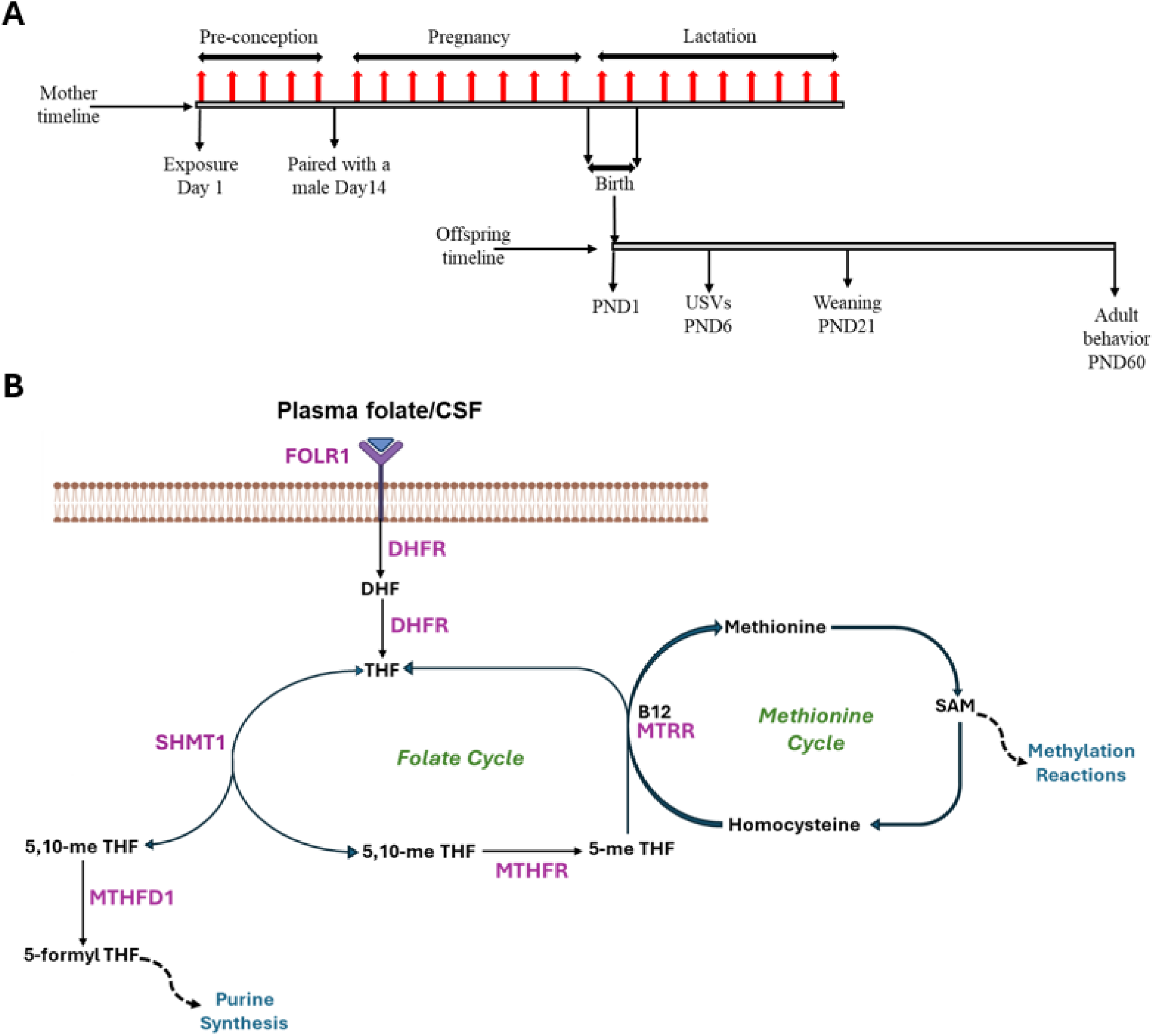
(A) Experimental design showing the experimental timeline of exposures in the dams and offspring. Exposures in the dams (indicated by red arrows) occurred across 2 weeks of preconception, then pregnancy and lactation. Time points of birth, weaning and behavior testing of the offspring are shown in post-natal days (PND). (B) The one-carbon metabolism pathway in cells showing the folate cycle and methionine cycle. Key enzymes: dihydrofolate reductase (DHFR); serine hydroxymethyltransferase 1 (SHMT1); methylene-tetrahydrofolate reductase (MTHFR); methionine synthase reductase (MTRR); methylene-tetrahydrofolate dehydrogenase 1 (MTHFD1); folate receptor α (FOLR1). Co-Factors and metabolites: dihydrofolate (DHF); tetrahydrofolate (THF); 5,10-methyltetrahydrofolate (5,10-me THF); 5-methyltetrahydrofolate (5-me THF); S-adenosylmethionine (SAM).

### Subject numbers and eliminations

Initially, our study included 80 dams (Control, N=24; DPE, N=26; DPE+F, N=28). The a priori elimination criteria for dams were (1) failure to consume 90% of the doses provided; (2) failure to give birth within 28 days to a litter containing at least one male and one female pup; and (3) fight-related injuries. One dam was eliminated for non-consumption. After being paired with a male, 23 dams were eliminated for not producing a balanced litter within the 28-day period. Five dams were eliminated for fight-related injuries. Some dams that produced a balanced litter had offspring of only one sex survive to adulthood; these litters were not eliminated. Eliminations did not differ between experimental groups. Following these eliminations, a total of 51 litters (Control, N=18; DPE, N=17; DPE+F, N=16) remained, which is sufficient to provide 80% power to detect an effect size of η^2^=0.168 prior to task-specific eliminations (described below). The offspring contained in these litters were divided among the experiments (Table S1). For operant conditioning only, another cohort of offspring were produced using identical methods (Control, N=10; DPE, N=13) and combined with the primary cohort.

### Experimental units

We followed the litter-based design standard for developmental exposure [2]. Because the experimental treatment was administered to the dam, the dam/litter was considered the experimental unit, with each litter counted as N=1. For each experiment, a representative animal was chosen from each litter as the test subject (Table S1), so the N for each experiment corresponded to both the number of animals and the number of litters represented. To reduce cross-interference between tests, separate animals from the same litter were used as subjects for behavior tests that involved foot shocks (fear conditioning and consolation) and food restriction (operant conditioning), and behaviorally naive animals were used for molecular assays. Experiments were conducted on one male or female offspring from each litter, except for protein assays, which were performed on tissues from one male and one female offspring from each litter with sex as a within-subjects factor.

### Blinding

Exposure doses were prepared as described below and blindly labeled by laboratory staff uninvolved in the experiments. Experimenters performing the exposures were blind to the identity of the groups. Cage numbers were randomly assigned to the offspring at weaning and were used throughout all subsequent tests. Experimenters remained blind to the identity of the groups until all data collection and analysis was complete.

### Exposure model decision points

We chose to model chronic low-dose ingestion of deltamethrin before, during, and after pregnancy; as well as folate supplementation before, during, and after pregnancy. The principal routes of exposure to pyrethroid pesticides in humans are ingestion and inhalation, with ingestion through food and hand transfer being the most common source of low-dose chronic exposure [53]. The type 1 pyrethroids (including deltamethrin) are rapidly metabolized, and deltamethrin has a half-life measured in hours. As such, in order to control for bioaccumulation of the exposure dose, we chose a phasic 3-day exposure interval. We also chose a single exposure dose of deltamethrin (3 mg/kg); this dose is well below the EPA-set benchmark dose of 14.5 mg/kg [43] and the previously observed developmental “no observable adverse effect level” (NOAEL) of 12 mg/kg [54], and is consistent with the maternal NOAEL of 2.5 - 3.3 mg/kg [55]. Importantly, this developmental exposure dose was sufficient to produce behavioral effects on mouse offspring in our previous studies [39]. For folate supplementation, the Centers for Disease Control (CDC) recommends daily supplementation with 400 µg folic acid before and during pregnancy [56]. The CDC also recommends that women at risk for a pregnancy affected by a neural tube defect take 4000 µg folic acid daily, or 10 times the normal amount. Based on the prairie vole laboratory diet, we estimated that the average prairie vole consumes approximately 0.5 mg/kg folate per day, so we selected a supplementation dose of 5 mg/kg. Finally, we chose to supplement folate using the bioactive form, methylfolate, to circumvent any potential effects of pesticide exposure on the conversion of folic acid into its bioactive form.

### Chemicals

Aliquots were made by dissolving deltamethrin (Sigma, St. Louis, MO) in acetone (Sigma) and 5-methyltetrahydrofolate di-sodium salt (Sigma) in water. Both solutions were suspended in cherry-flavored veterinary compounding syrup (Humco, Texarkana, TX) and left overnight in the fume hood for the acetone to evaporate. Control aliquots were prepared identically by mixing the same volumes of acetone and water into veterinary compounding syrup and allowing the acetone to evaporate. The resulting mixtures were aliquoted in borosilicate glass vials and stored at −80°C until the day of use. On each exposure day, aliquots were thawed and vortexed immediately prior to use.

### Deltamethrin exposures

Prior to the experiment, potential prairie vole dams were pre-exposed to unmodified cherry-flavored veterinary compounding syrup daily for three days. Animals that did not consume the syrup for at least two consecutive days were not used for the study. On the day of each experimental exposure, females were removed from the home cage and weighed, and a weight-adjusted volume of syrup was drawn into a 1mL open-tipped syringe. The dams were lightly restrained and introduced to the tip of the loaded syringe. The animals would voluntarily consume the syrup by licking it off the open tip of the syringe while the appropriate volume was being dispensed. Dams were orally exposed to cherry syrup containing deltamethrin (3mg/kg), vehicle, or deltamethrin (3 mg/kg) with methylfolate (5mg/kg) once every three days. Exposures started two weeks prior to being paired with a male, and continued throughout preconception, pregnancy, and the lactation period. All feedings were strictly done between 10 AM and 2 PM. The offspring were only exposed indirectly through the dam, with exposures terminating at weaning.

### Behavioral batteries

A variety of complementary behavioral assays were chosen to represent five key domains relevant to NDDs across the lifespan: the social, communication, cognitive, locomotor, and repetitive behavior domains (Table S2) [50]. Many of the assays were selected to replicate the effects of DPE in mice [39]; assays replicated from the mouse used one-tailed statistical tests when available, since the only question of scientific interest was whether the previously observed phenotype was present. To reduce the impact of repeated testing stress as a confounding factor, the assays were organized into a fixed-order behavioral battery in ascending order of stress, with the least stressful test performed first. The order of testing (as described below) was ultrasonic vocalizations, observation, marble burying, 24-hour movement, partner preference, and then consoling behavior. Assays involving foot shock (fear conditioning) and food restriction (operant conditioning) were performed on separate animals from the same litter. One male or female vole per litter was used for each assay, except for ultrasonic vocalizations, where the sex of pups (postnatal day 6-7) was not identified.

### Separation-induced ultrasonic vocalizations (USVs)

We recorded separation-induced USVs at postnatal day (PND) 6-7 from a single offspring from each litter as previously described for mouse [39]. One data point was removed due to a corrupted audio file (Control, N=17; DPE, N=17; DPE+F, N=16). Male and female prairie vole pups are indistinguishable at PND 6-7, so the sex of pups was not determined. Recordings were made inside a sound-attenuating recording booth (ROOM, Brooklyn, NY) underneath a Noldus microphone attached to Noldus Ultravox software (Noldus, Wageningen, The Netherlands). The resulting audio files were analyzed for call number and seven additional acoustic parameters using the DeepSqueak v3.0 software package [57] for MatLab (Version R2023a, Mathworks, Natick, MA), using the general purpose network with default settings. Acoustic parameters included call number, duration, principal frequency, peak frequency, delta frequency, slope, sinuosity, and tonality (Table S2). Call number was our measure of interest and was analyzed using a two-factor ANOVA (factors of group and sex); post hoc one-tailed t-tests were used as this measure was replicated from mouse. Exploratory two-factor ANOVAs were used to analyze all other acoustic parameters.

### Novel cage observation

The novel cage observation test was performed and analyzed for spontaneous repetitive behaviors as previously described [39]. Digital recordings were scored automatically using the Noldus Ethovision software (Ethovision 16 XT, Noldus). Self-grooming and rearing were our outcomes of interest. Data from self-grooming and rearing were analyzed using two-factor ANOVAs (factors of group and sex); post hoc one-tailed t-tests were used as this measure was replicated from mouse.

### Marble burying

The marble burying test was performed as previously described [39]. Our primary measure of interest was number of marbles buried, which was scored by agreement between three independent, blinded raters. We compared marbles buried between treatment groups using a two-factor ANOVA (factors of group and sex); post hoc one-tailed t-tests were used as this measure was replicated from mouse.

### 24-hour movement and locomotor activity tests

Movement was measured using a combined test for locomotor activity and 24-hour movement patterns. Each adult offspring was transferred into an 18”x18”x18” open field cage inside a sound-attenuating recording booth. The animals were provided with food and water ad libitum and their total movement was recorded for a 25-hour period. Start time was counterbalanced throughout the day. The recording booths were equipped with a hygrometer-thermometer (Thermopro, Amazon.com, Seattle, WA) and hue-changing smart lights (Philips, Amsterdam, The Netherlands) programmed to maintain the day-night cycle by shifting between white light and low-level red light. All recordings were scored using Ethovision software (Ethovision 16 XT, Noldus). The first hour of the test was considered novelty-induced locomotor activity, and the next 24 hours were considered 24-hour movement. Novelty-induced locomotor activity was compared between treatment groups using a two-factor ANOVA (factors of group and sex) with post hoc one-tailed t-tests, as this measure was replicated from mouse [30]. 24-hour movement data was organized into 1-hour time bins based on Zeitgeber (ZT) time, which is the number of hours since lights-on. Due to a technical problem, a block of time was missing from a small subset of subjects, so 24-hour movement was compared between groups using a linear mixed model. Missing data points were not imputed because they were sequential. Post hoc one-way ANOVAs were used to test for the effect of group at each time point.

### Partner preference test

The partner preference test was performed as previously described [58, 59]. Male and female subjects were separated from their same-sex cagemates and paired with a sexually naïve opposite-sex mating partner from the same experimental group for 24 hours of cohabitation. One member of the pair was designated as the “subject” for all subsequent testing. At the end of cohabitation, each subject’s partner was removed from the home cage and a zip-tie “collar” attached by a fishing chain “leash” to a magnetic anchor was fitted around their neck. The collar and anchor was then used to restrict the partner’s movement to one chamber of a three-chambered testing cage. Another subject’s partner was confined to the opposite end of the same cage, and this cage then served as the test chamber for both subjects, with each partner serving as the “stranger” for the other subject. Partners were allowed 15 minutes to habituate before one of the two test subjects was introduced to the center cage. The test subject was allowed to freely move through all three chambers for 3 hours, and digital video was recorded. At the end of 3 hours, the partners were briefly removed from the testing cage, the cage was cleaned and lined with fresh bedding, the partners were returned, and the second subject was tested for another 3 hours. Food and water was provided throughout the test. At the end of testing, subjects and partners were reunited and remained paired for all subsequent experiments. Seven subjects were eliminated when the test was stopped due to fighting (Control, N=9; DPE, N=8; DPE+F, N=12). Eliminations did not differ statistically between experimental groups.

Top-down video recordings of partner preference tests were analyzed using CleverSys Topscan software (CleverSys, Reston, VA) as described previously [59]. Briefly, each test chamber was divided into three zones: two social zones on either end and a middle zone in between. Huddling was defined as immobile social contact with either the partner or the stranger within these social zones, and the duration of time spent huddling with the partner and stranger was measured for each subject. Because individuals in our colony vary in coat color, the contrast values used to track subjects against the background ranged from 15-20, and the maximum pixel value used to track subjects ranged from 40-125 (0 is black, 255 is white). Huddling time with the partner and stranger was transformed to ranks and compared between groups using a repeated measures ANOVA (within-subjects factor of partner; between-subjects factors of group and sex) followed by post hoc two-tailed paired t-tests.

### Consoling behavior

The consoling behavior test was performed as previously described [45] using male-female pairs that were previously paired for partner preference testing. One pair was eliminated due to an unrelated injury (Control, N=12; DPE, N=12; DPE+F, N=11). Reunion sessions were manually coded by agreement between two blind raters using Noldus Observer (Noldus, Waneningen, The Netherlands), with observer allogrooming (i.e. the consoling response) as the primary measure of interest. Consoling response was converted to ranks and compared between groups using a two-factor ANOVA (factors of group and sex) with post hoc two-tailed t-tests.

### Classical fear conditioning

Classic fear conditioning was performed in a conditioning chamber (Coulbourne Instruments, Holliston, MA) as previously described [39] concurrently with consoling behavior, with the animal designated as the “demonstrator” undergoing conditioning. One pair was eliminated due to an unrelated injury (Control, N=12; DPE, N=12; DPE+F, N=11). As previously, a 3-day protocol was used with fear acquisition (5 tones), fear recall (5 tones), and fear extinction (30 tones). Freezing behavior throughout the test was measured using FreezeFrame5 software (Actimetrics, Wilmette, IL). Our primary measure of interest was freezing during the first 5 tones of day 2 (fear recall), with freezing during tones on the first day (fear acquisition) and freezing during all 30 tones on days 2 and 3 (fear extinction) as secondary measures. Freezing was compared between groups using a repeated measures ANOVA (within-subjects factor of tone; between-subjects factors of group and sex) with post hoc t-tests.

### Operant conditioning

Adult offspring from a subset of litters, plus additional subjects from a separate cohort (Table S1), were trained to nose poke for food using the previously described protocol [39]. Briefly, voles were subjected to food restriction consisting of daily weighed food portions and targeting a body weight of 90-95% of their free-feeding weight. Eight voles fell below 85% of their free-feeding weight for 2 days in a row and were eliminated, leaving a total of 55 subjects (Control, N=20; DPE, N=23; DPE+F, N=12). Eliminations did not differ significantly between groups. For 70 minutes per day for 10 consecutive days, voles were placed in Panlab 5-choice operant conditioning chambers (Harvard Apparatus, Holliston, MA) with two active nose poke apertures and a liquid dispenser containing a 50% nutritional shake solution (Original Strawberry Ensure, Abbott, Chicago, IL) diluted with tap water. Voles could earn up to 30 drops (50 µL each) per aperture per day on a fixed ratio of one (FR1) schedule, for a maximum of 3 mL/day. For each daily training session, voles were removed after earning 60 drops or after the total test time elapsed. Voles were considered to have passed the acquisition criterion if they earned 30 drops from each aperture on any combination of days. As previously in mice [39], acquisition criterion was the primary outcome of interest, and was compared between groups using a chi-squared table test with post hoc chi-squared tests. Data on the daily “time to completion” was only collected from one cohort of subjects (DPE, N=12; Control, N=12; DPE+F, N=12) and these data were used to calculate the nose poke rate, from which a 10-day learning curve was constructed. The 10-day learning curve was compared between groups using a repeated measures ANOVA with post hoc two-tailed t-tests. The slope of the learning curve for this subset of subjects was calculated using the SLOPE function in Excel (version 2405, Microsoft, Redmond, WA) and compared between groups using a Kruskall-Wallis test with post hoc one-tailed U tests.

### Integrated behavioral Z-scoring

The combined behavioral phenotype was quantified using the integrated behavioral Z-scoring method described by Guilloux et al [60, 61] using a procedure that weighted the five behavioral domains equally. Continuous outcome measures from each domain were selected (Table S2). Outcomes were selected post hoc based on the presence of a statistical difference, making this measure descriptive. The primary outcome measures for operant conditioning and 24-hour mobility were not single continuous variables, so secondary outcome measures (slope of the learning curve, and movement during Zeitgeber time 7-8, respectively) were included instead. Z-scores for the selected outcomes were calculated relative to the mean and standard deviation of the control group, first at the subject level, then averaged within each litter, then averaged within each of the five domains (Supplemental Data File 1B). Domain Z-scores were adjusted for directionality to reflect deficits as positive values (i.e. communication, cognitive, and social domain scores were inverted) and then averaged between domains to obtain a single integrated behavioral Z-score per litter. Integrated Z-scores were compared between groups using a one-way ANOVA with post hoc t-tests.

### Dissections, tissue collection, and sample preparation

Samples from behaviorally naïve offspring were used for all molecular assays. All offspring were euthanized via carbon dioxide euthanasia followed by decapitation. Trunk blood was collected in EDTA tubes (Eppendorf, Hamburg, Germany) on ice, centrifuged at 8000 RPM for 10 minutes, and the plasma fraction was collected and stored at −80°C. Brain, liver, and fecal samples were collected, rapidly frozen on dry ice, and stored at −80°C. Whole brain tissue lysates were later prepared by suspending brain samples in CelLytic MT cell lysis buffer (Sigma) freshly supplemented with a protease inhibitor cocktail powder (P2714-1BTL, Sigma) and homogenizing the sample in a bead beater (TissueLyser LT, QIAGEN, Ann Arbor, MI) for 3 minutes at 50 Hz. The homogenized mixture was centrifuged for 20 mins at 13,000 rpm, and the supernatant was collected and passed through a QIAshredder (QIAGEN) before being checked for concentration using BCA assay (Thermo Scientific, Waltham, MA). The resulting tissue lysates were aliquoted for Western blot and chemiluminescence assay (below).

### Chemiluminescence assay

Blood plasma samples and whole brain tissue lysates were prepared as described above. A randomly selected subset of plasma samples and brain lysates (N=14 per sample type per group) were analyzed for Vitamin B9/Folic acid using a commercially available chemiluminescent immunoassay (CLIA) kit (Abbexa, Cambridge, UK) that reports the combined concentration of all molecular forms of folate detected. Experimental procedure was followed in accordance with kit user-manual. The resulting optical densities were measured and recorded using Cytation 5 (Agilent BioTek, Santa Clara, CA). Assay controls were used to construct a standard curve, and sample optical density was converted to concentration, which was our primary outcome measure. Data was compared between treatment groups using two-factor ANOVAs (factors of group and sex) with post hoc two-tailed t-tests.

### Antibodies

We selected six critical proteins from the one-carbon pathway as targets for analysis (Fig. 1B): folate receptor α (FOLR1), methylene-tetrahydrofolate reductase (MTHFR), serine hydroxymethyltransferase 1 (SHMT1), dihydrofolate reductase (DHFR), methionine synthase reductase (MTRR), and methylene-tetrahydrofolate dehydrogenase 1 (MTHFD1). Vinculin, a stable ubiquitously expressed cytoskeleton protein with expected bands not overlapping with our selected antibodies, was used as a loading control. Primary antibodies used for Western blots included rabbit anti-FOLR1 (ab221543, 1:2000 dilution, Abcam, Cambridge, UK), rabbit anti-MTHFR (ab203786, 1:2000 dilution, Abcam), rabbit anti-SHMT1 (80715S, 1:2000 dilution, Cel Signaling Technology, Danvers, MA), rabbit anti-DHFR (ab288373, 1:2000 dilution, Abcam), rabbit anti-MTRR (NBP2-93732, 1:2500 dilution, Novus Biologicals, Centennial, CO), mouse anti-MTHFD1 (ab70203, 1:500 dilution, Abcam), and mouse anti-vinculin 7F9 (sc-73614, 1:500 dilution, Santa Cruz Biotechnology, Santa Cruz, CA). Secondary antibodies used included IRDye 680RD Donkey anti-Rabbit (926-68073, 1:10,000 dilution, LI-COR, Lincoln, NE) and IRDye 800CW Donkey anti-Mouse (926-32212, 1:10,000 dilution, LI-COR).

### Western blots

Protein immunoreactivity in a Western blot was measured as a proxy for protein expression. Whole brain tissue lysates were mixed with SDS loading buffer (SDS Sample Loading Buffer 4X-21420018, bioWORLD, Dublin, OH) and boiled at 85°C for 5 mins. 20µg of protein was used for polyacrylamide gel electrophoresis (10% Mini-PROTEAN TGX Precast 15-well Protein Gels, Biorad, Hercules, CA) followed by transfer to a PVDF membrane (Merck KgaA, Darmstadt, Germany) and saturation in Odyssey Blocking Buffer (LI-COR, Lincoln, NE) for 5 mins. Membranes were then incubated with primary antibody overnight at 4°C, followed by wash steps and incubation with secondary antibody for 1 hour at room temperature. Membranes were then washed in Tris-buffered saline with 0.1% Tween 20 and imaged using LI-COR Odyssey Blot Imager (LI-COR). Band density was quantified using ImageJ software to measure the optical density, subtract the background immediately surrounding the band, then calculate the ratio to the band density (minus background) of the loading control. We used this unitless ratio of relative protein expression as the primary measure of interest. We processed samples from one male and one female offspring from each litter, and due to the litter-based design, sex was used as a within-subjects factor for analysis. Only litters for which one male and one female sample were available were included in the final analysis (Control, N=13; DPE, N=11; DPE+F, N=13). Data was analyzed using repeated measures ANOVAs (within-subjects factor of sex; between-subjects factor of group) with post hoc two-tailed t-tests.

### Statistics

SPSS version 29 (IBM Corp., Armonk, NY) was used to perform group comparisons using parametric and non-parametric statistical tests. GraphPad Prism version 10 (GraphPad, La Jolla, CA) was used for linear mixed models and for graph generation. Experimental data that were both parametrically distributed (by Kolmogorov-Smirnov test) and had homogeneity of variance (by Levine’s test or Mauchly’s sphericity) were analyzed using parametric statistics. Data that failed either criterion were either transformed to fractional ranks or analyzed using nonparametric statistics. Data containing repeated measures with missing data points were analyzed using linear mixed models instead. Omnibus tests (ANOVA, Kruskal-Wallis, linear mixed model, chi-squared table) were considered the primary analysis; uncorrected post hoc tests (t-test, U-test, chi-squared) were used descriptively to describe the differences between groups detected by the primary analysis. Effect sizes (in partial eta squared or Cohen’s d) were calculated and reported for all statistical tests. Specific assays replicated from the mouse (Table S2) used one-tailed statistical tests when available, since the only question of scientific interest was whether the previously observed phenotype was present. Factors and post hoc tests used for each statistical comparison are mentioned in the Methods for each assay above.

### Data management

All project-related files were stored and shared between investigators within and between Universities using Dropbox Business cloud storage (Dropbox Inc, San Francisco, CA).

## RESULTS

### Developmental exposure

Prairie vole dams (N=51) were exposed to experimental treatment every 3 days during pre-conception, pregnancy, and lactation (Fig. 1A). Experimental treatments given to the dams consisted of orally consumed veterinary syrup containing vehicle, deltamethrin, or deltamethrin plus the folate vitamer methylfolate. There were no differences between groups in average litter size, birth weight, or number of days between pairing and birth (ANOVAs, p>0.05). The resulting offspring were considered the subjects of subsequent experiments, with N=17 litters developmentally exposed to vehicle (Control), N=18 litters developmentally exposed to pesticide (DPE), and N=16 litters developmentally exposed to pesticide plus methylfolate (DPE+F).

### Social domain

Prairie vole offspring were tested at adulthood on their species-specific complex social behaviors, as measures relevant to the social domain (Fig. 2A-B). In the partner preference test, all exposure groups preferred their partner to an opposite-sex stranger after 24 hours of cohabitation (ANOVA, main effect of partner, F(1,26)=47.4, η^2^ =0.65, p<0.001) with no differences in preference between groups (ANOVA, main effect of group, F(2,26)=0.85, η^2^ =0.06, p=0.44) (Fig. 2A). However, in the consolation test, exposure groups differed in their response to distressed partners (ANOVA, F(2,32)=3.3, η^2^ =0.17, p=0.049), with DPE voles showing a reduced consoling response compared to both controls (t-test, t(22)=0.031, d=0.94, p=0.031) and DPE+F voles (t-test, t(21)=2.1, d=0.87, p=0.050) (Fig. 2B). This result demonstrates that DPE induced a deficit in complex social behavior that was prevented with maternal methylfolate supplementation.

**Figure 2.**
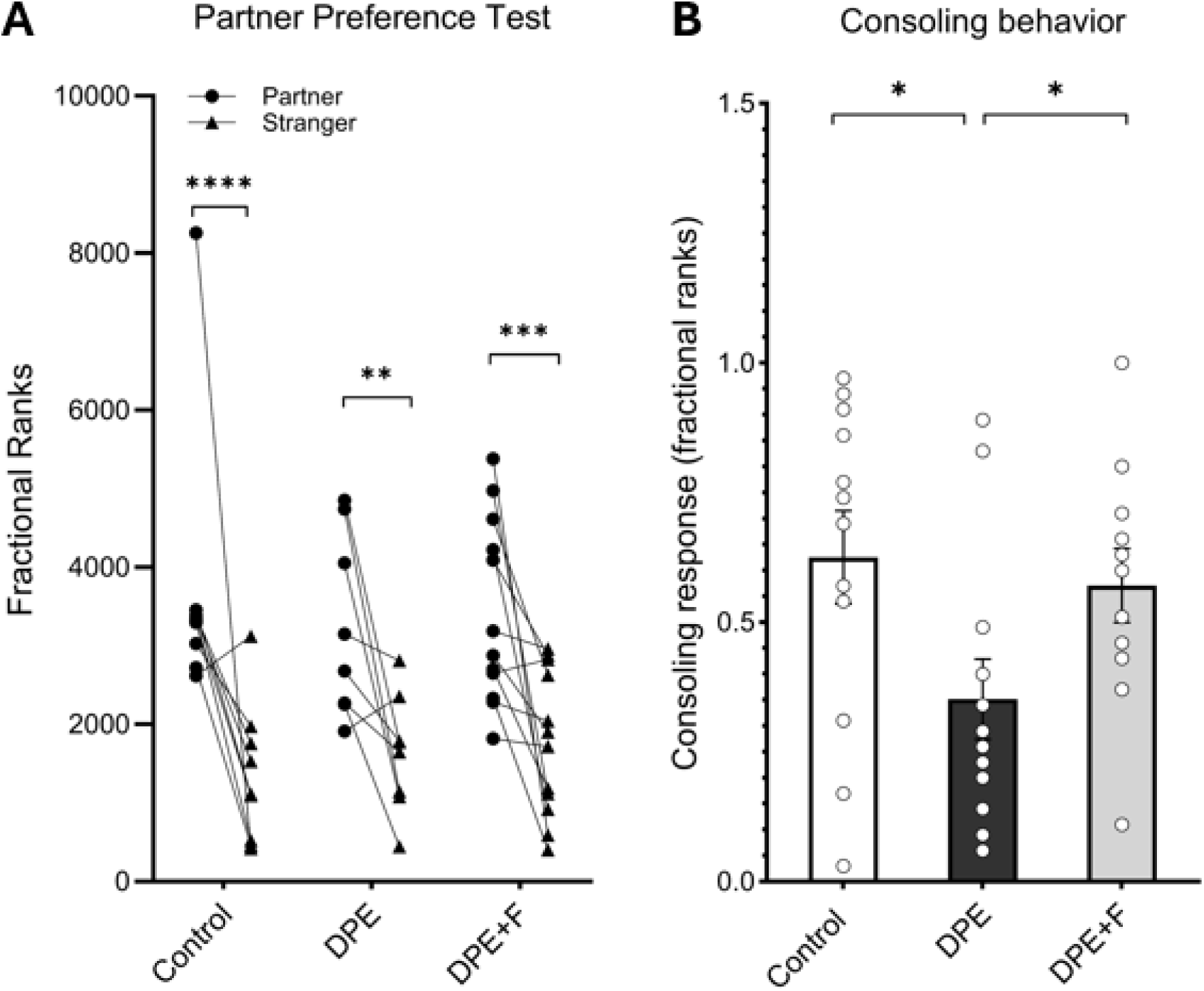
Species-specific social behaviors in prairie voles. (A) Prairie voles in all experimental groups (Control, N=9; DPE, N=8; DPE+F, N=12) showed a significant preference for their partner over a stranger after 24 hours of cohabitation. (B) DPE voles (N=12) showed reduced consoling responses toward their distressed partners relative to controls (N=12) and DPE+F voles (N=11). Error bars are standard error. * p < 0.05 ** p < 0.01 *** p < 0.005 **** p < 0.001

### Communication domain

Prairie vole offspring were tested for separation-induced vocalizations at 6-7 days of age, as a measure relevant to the communication domain (Fig. 3A-B). The total number of prairie vole pup vocalizations produced differed between exposure groups (ANOVA: F(2,47)=5.5, η^2^ =0.19, p=0.007), with DPE pups producing fewer vocalizations than either control pups (t-test, t(32)=2.7, d=0.91, p=0.006) or DPE+F pups (t-test, t(31)=3.0, d=1.0, p=0.003) (Fig. 3A). As an additional exploratory analysis, vocalizations were assessed for variation in seven acoustic parameters (Table S2). There were group differences in the duration of ultrasonic vocalizations (ANOVA: F(2,47)=4.5, η^2^ =0.16, p=0.017; Fig. 3B) but not for other acoustic parameters like peak frequency, sinuosity or tonality (ANOVAs: p > 0.05). Specifically, DPE pups produced shorter vocalizations than both control pups (t-test: t(32)=2.6, d=0.91, p= 0.013) and DPE+F pups (t-test: t(31)=2.6, d=0.90, p=0.016). This pattern demonstrates that DPE induced a deficit in the amount of calling that was prevented with maternal methylfolate supplementation.

**Figure 3.**
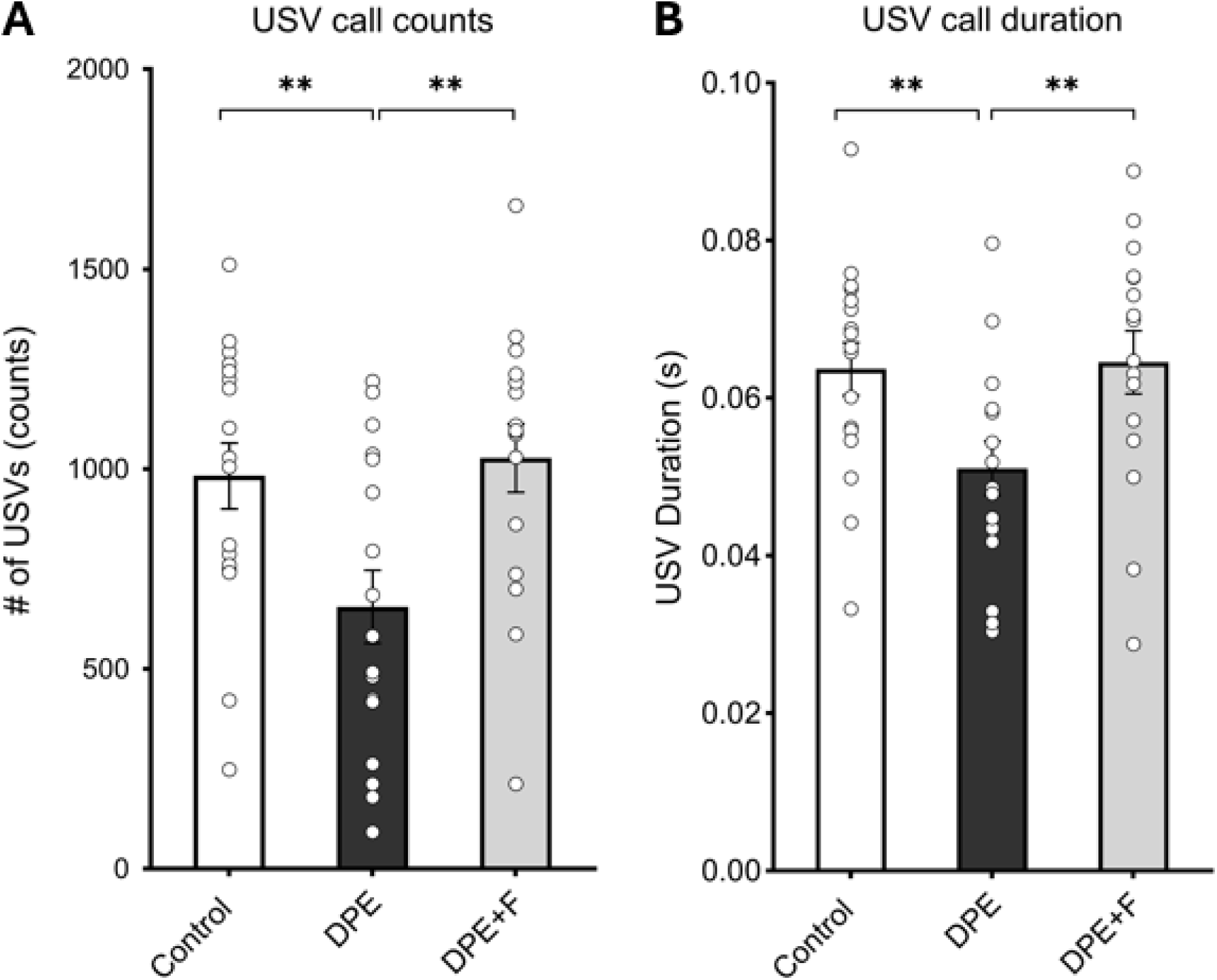
Separation-induced ultrasonic vocalizations (USVs) in 1-week old pups. (A) DPE pups (N=17) produced fewer USVs than controls (N=17), an effect which was prevented in DPE+F pups (N=16). (B) DPE voles had shorter average call durations than controls and DPE+F pups. Error bars are standard error. * p < 0.05 ** p < 0.01 *** p < 0.005 **** p < 0.001

### Cognitive domain

Adult prairie vole offspring were tested for learning-related outcomes, as measures relevant to the cognitive domain (Figs. 4A-C, S1A-B). Offspring underwent operant conditioning under food restriction, consisting of nose pokes reinforced with food reward on an FR1 schedule. There were no differences between groups in adult body weight, food consumption by weight, or eliminations (ANOVAs, p>0.05). At the end of 10 days of training, exposure groups differed in the proportion of subjects acquiring the operant response (χ^2^ table test, χ^2^=7.7, p=0.021), with fewer DPE voles acquiring the response than controls (χ^2^ test, χ^2^=6.6, p=0.010) or DPE+F voles (χ^2^ test, χ^2^=4.7, p=0.030) (Fig. 4A). The nose poke rate per day across 10 days differed between exposure groups (ANOVA, time x group interaction, F(18,297)=2.6, p=<0.001), with DPE voles differing from DPE+F voles in the latter half of training (t-tests: Day 7, t(22)=2.4, d=0.98, p=0.026; Day 8, t(22)=2.2, d=0.90, p=0.038) (Fig. 4B). The slope of the 10-day learning curve also differed between exposure groups (Kruskal-Wallis test, H=6.6, p=0.037), with DPE voles having a slope consistent with zero, which was lower than both control voles (U test, U=39.5, p=0.03) and DPE+F voles (U test, U=30.00, p=0.0.006) (Fig. 4C). In the fear conditioning experiment, all exposure groups acquired the fear response (ANOVA, main effect of time, F(4,112)=27.6, η^2^=0.50, p<0.001) and extinguished the fear response (ANOVA, main effect of time, F(29,812)=8.3, η^2^=0.23, p<0.001) with no differences detected between groups on measures of fear acquisition (ANOVA, group x time interaction, F(8,112)=1.2, η^2^=0.077, p=0.33), fear recall (ANOVA, main effect of group, F(2,28)=1.1, η^2^=0.071, p=0.36), or fear extinction (ANOVA, group x time interaction, F(58,812)=1.1, η^2^=0.0.072, p=0.32) (Figs. S1A-C). This pattern demonstrates that DPE induced a deficit in the cognitive domain that was prevented with maternal methylfolate supplementation.

**Figure 4.**
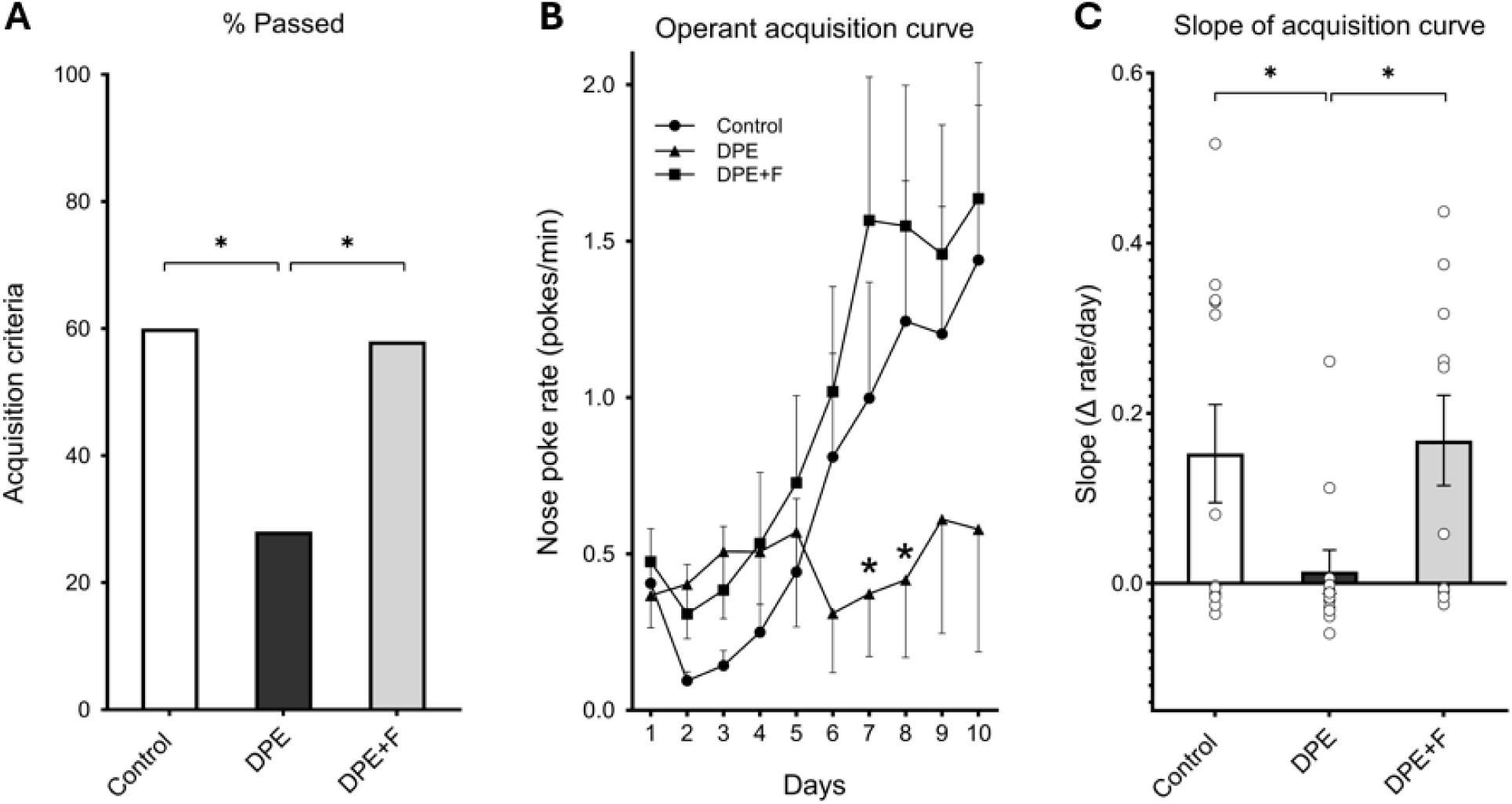
Operant conditioning of a nose poke response to acquire food. (A) Across two independent cohorts, DPE voles (N=20) were less likely to meet the criteria for acquiring the operant response after 10 days relative to both controls (N=23) and DPE+F voles (N=12). (B) DPE voles (N=12) did not increase their nose poke rate over 10 days of training relative to both control (N=12) and DPE+F voles (N=12). (C) The slope of the acquisition curve was significantly lower in DPE voles (N=12) relative to control (N=12) and DPE+F voles (N=12). Error bars are standard error. * p < 0.05 ** p < 0.01 *** p < 0.005 **** p < 0.001

### Locomotor domain

Adult offspring were tested for hyperactivity and 24-hour movement patterns, as measures relevant to the locomotor domain (Fig. 5A-C). Total movement in a novel environment differed between exposure groups (ANOVA main effect of group, F(2,41)=7.3, η^2^=0.26, p=0.002), with DPE voles showing greater total movement than both control voles (t-test, t(31)=2.7, d=0.93, p=0.006) or DPE+F voles (t-test, t(28)=1.9, d=0.68, p=0.037) (Fig. 5A). Hyperactivity seemed to be most pronounced in females (ANOVA group x sex interaction, F(2,41)=3.9, η^2^=0.16, p=0.029), with DPE females showing hyperactivity relative to controls (t-test, t(12)=2.7, d=1.5, p=0.010) and DPE+F females (t-test, t(10)=2.2, d=1.3, p=0.025). Exposure groups also differed in their 24-hour movement patterns (mixed model, F(46,963)=1.5, p=0.0258), with DPE voles showing increased movement in the hours leading up to, and certain portions of, the dark cycle (Fig. 5B). This pattern demonstrates that DPE induced multiple deficits in the movement domain that were prevented by maternal methylfolate supplementation.

**Figure 5.**
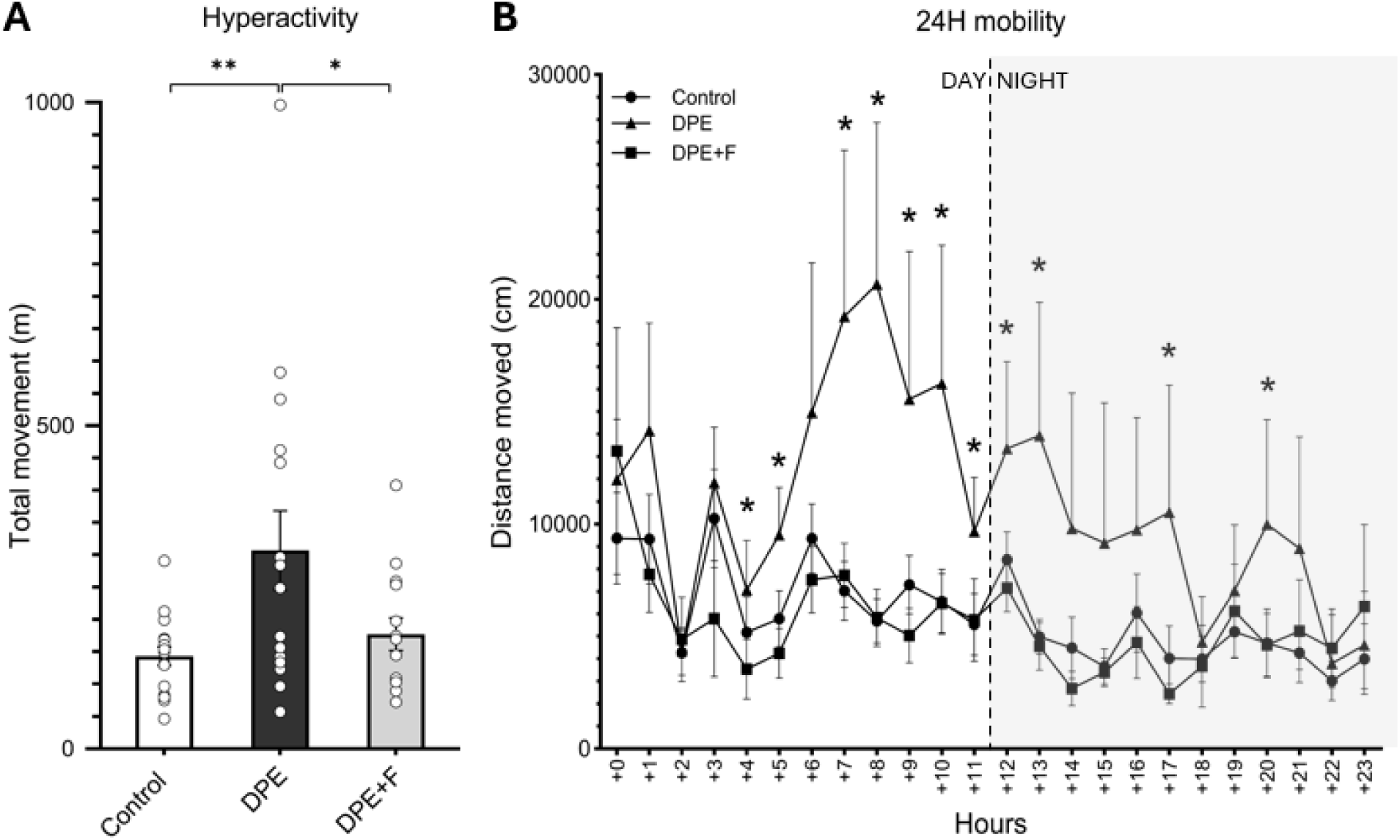
Locomotor activity. (A) DPE voles (N=16) were hyperactive relative to controls (N=17) and DPE+F voles (N=14). (B) Prairie voles have both circadian and ultradian rhythms, with 3-4 hour ultradian movement cycles that are more pronounced in the light than in the dark. DPE voles showed disrupted 24-hour movement patterns, with an increase in activity in the latter half of the light cycle and portions of the dark cycle. Error bars are standard error. * p < 0.05 ** p < 0.01 *** p < 0.005 **** p < 0.001

### Repetitive behavior domain

Adult offspring were tested for a variety of repetitive behaviors (Fig. 6A-C). In the marble burying test, there were differences between exposure groups in the number of marbles buried (ANOVA, F(2,41)=6.1, η^2^=0.23, p=0.005), with the DPE+F voles burying more marbles relative to both controls (t-test, t(29)=3.3, d=1.2, p=0.001) and DPE voles (t-test, t(28)=2.7, d=1.0, p=0.005) (Fig. 6A). In the observation test, there were differences between exposure groups in repetitive rearing behavior (ANOVA, main effect of group, F(2,41)=15.4, η^2^=0.43, p=<0.001), with DPE voles rearing more than control voles (t-test, t(31)=2.5, d=0.88, p=0.009) and DPE+F voles rearing more than both other groups (t-tests: Control x DPE+F, t(29)=5.6, d=2.0, p<0.001; DPE x DPE+F, t(28)=2.1, d=0.78, p=0.021) (Fig. 6B). Among DPE voles, the effect of DPE on rearing was more pronounced among females (ANOVA, sex x group interaction, F(2,41)=5.2, η^2^=0.20, p=0.010; paired t-test, male x female, t(14)=3.6, d=1.9, p=0.002). No differences in self-grooming between groups were detected (ANOVA, F(2,44)=1.4, η^2^=0.061, p=0.2517), but the same pattern was evident with a medium effect size (Fig. 6C). This pattern demonstrates that DPE induced a deficit in the repetitive and restrictive behavior domain that may have been exacerbated by maternal methylfolate supplementation.

**Figure 6.**
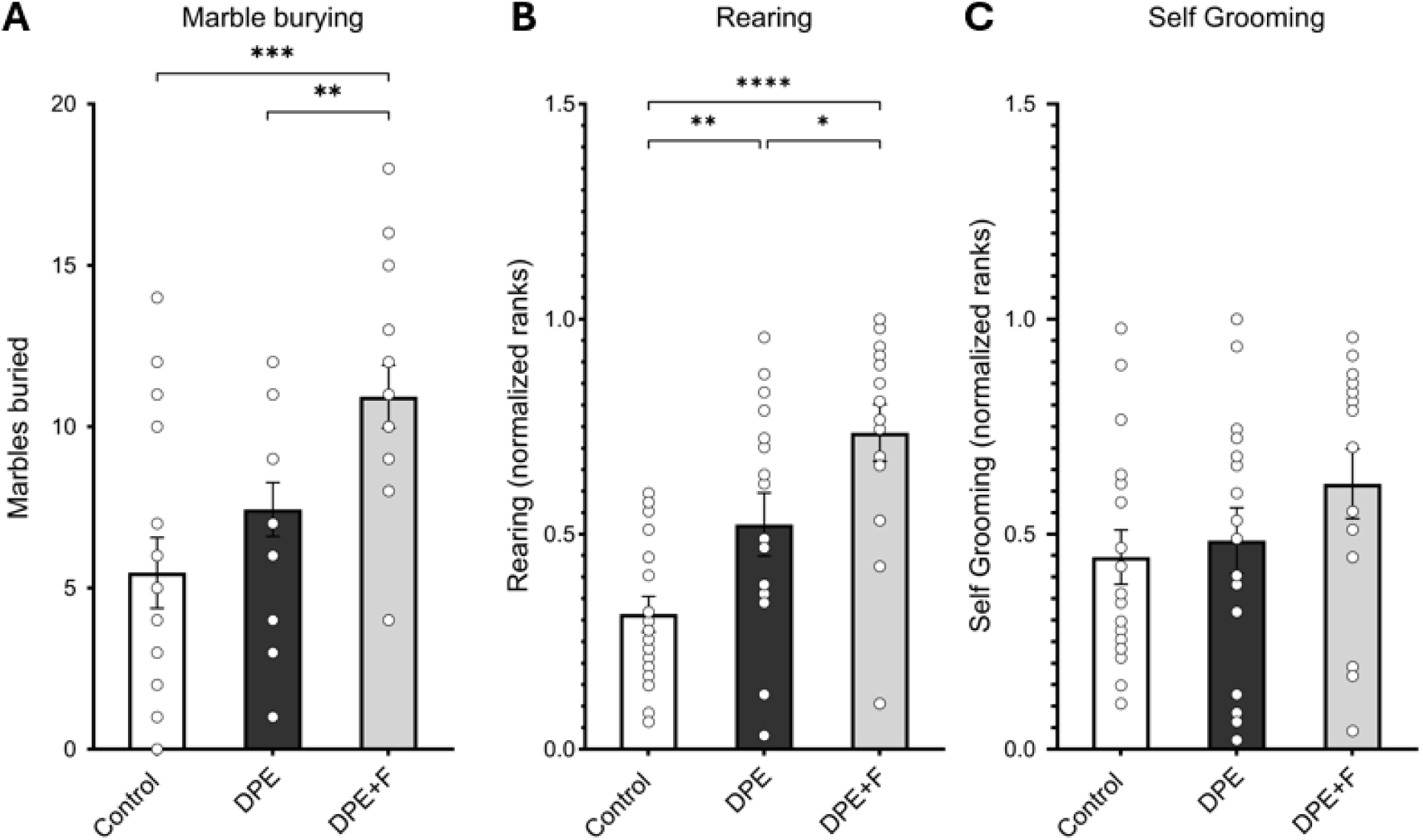
Repetitive behaviors. (A) DPE+F voles (N=14) buried more marbles as a consequence of repetitive digging compared to both control (N=17) and DPE voles (N=16). (B) When placed in a novel cage, DPE voles (N=16) showed more repetitive rearing than controls (N=17), and DPE+F voles (N=14) showed more than both DPE and controls. (C) There were no significant differences in repetitive self-grooming between groups. Error bars are standard error. * p < 0.05 ** p < 0.01 *** p < 0.005 **** p < 0.001

### Combined behavioral phenotype

To describe the combined behavioral phenotype across all five behavioral domains, integrated behavioral Z-scores were calculated using a method that weighted the domains equally (Fig. 7, Table S2, Supplemental Data File 1B). Comparing this integrated Z-score across groups revealed group differences (ANOVA, F(2,48)=6.6, η^2^=0.22, p=0.003) showing that DPE induced a consistent and strong combined phenotype (Z=1.3σ on average) as compared to controls (t-test, Control x DPE, t(33)=2.9, d=0.98, p=0.007), and that maternal methylfolate supplementation reduced this DPE phenotype (t-test, DPE x DPE+F, t(32)=2.4, d=0.82, p=0.024) to parity with controls (t-test, Control x DPE+F, t(31)=1.0, d=0.37, p=0.31).

**Figure 7.**
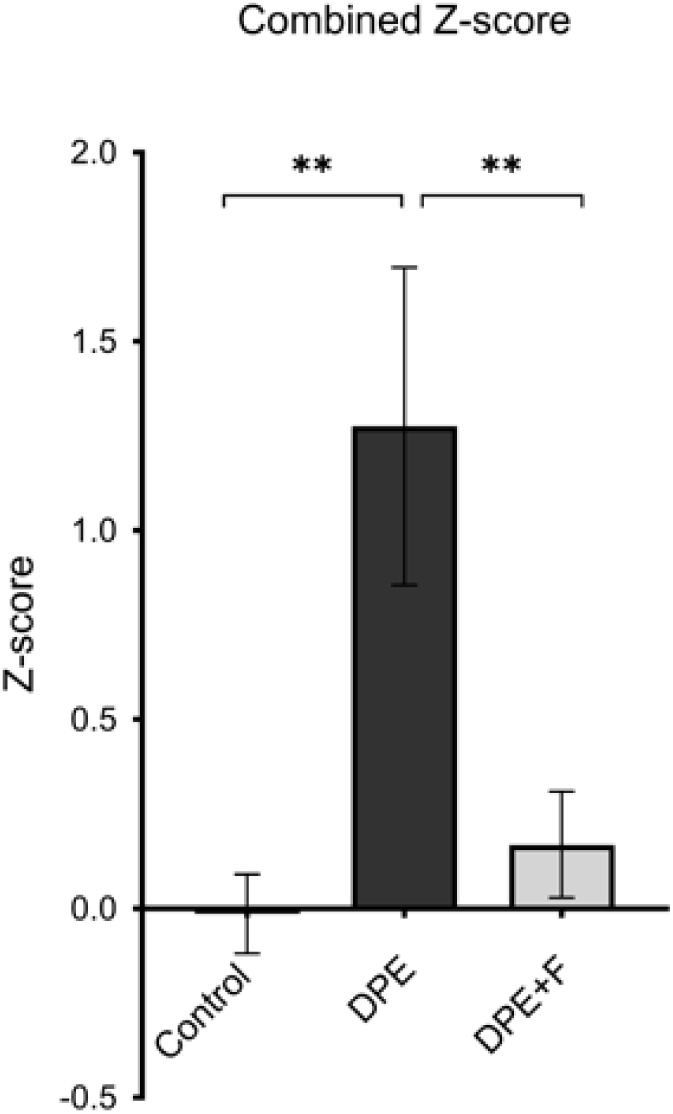
The combined Z-scores across all behavioral battery testing showed that DPE voles (N=18) exhibited a consistent change in their behavioral phenotype relative to controls (N=17), and that maternal folate supplementation (N=16) largely prevented the emergence of this phenotype. Error bars are standard error. * p < 0.05 ** p < 0.01 *** p < 0.005 **** p < 0.001

### Plasma and cerebral folate

Folate concentration was measured in plasma and brain lysate collected from adult offspring (Fig. 8A-B). Plasma folate concentrations differed across all three exposure groups (ANOVA, F(2,36)=15.2, η^2^=0.46, p=<0.001), with plasma folate elevated in DPE voles (t-test, t(26)=3.6, d=1.3, p=0.001) but relatively normalized in DPE+F voles (t-tests: DPE x DPE+F, t(26)=5.2, d=2.0, p=<0.001; Control x DPE+F, t(26)=2.0, d=0.75, p=0.06) (Fig. 8A). No differences in cerebral folate concentrations were detected (ANOVA, F(2,36)=1.5, η^2^=0.075, p=0.25), but the same pattern was evident with a medium effect size due to increased variability (Fig. 8B). No effects of sex were detected on either measure (ANOVAs, p>0.05).

**Figure 8.**
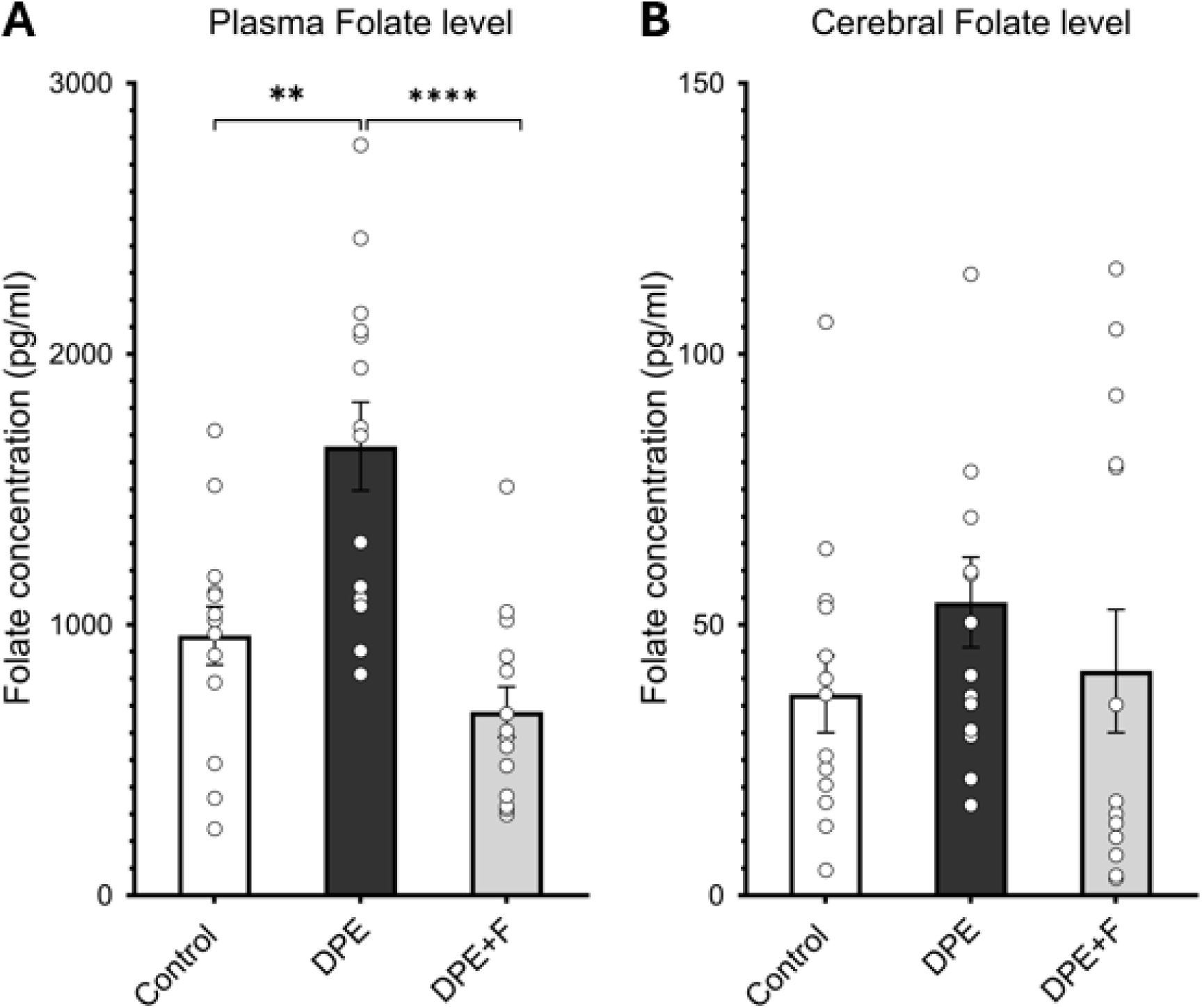
Folate concentrations in blood and brain as measured by a chemiluminescence assay that produces a combined response to all forms of folate detected. (A) DPE voles (N=14) had increased plasma folate concentrations relative to controls (N=14), and this increase was prevented in DPE+F voles (N=14). (B) There were no significant differences in cerebral folate concentrations between groups. Error bars are standard error. * p < 0.05 ** p < 0.01 *** p < 0.005 **** p < 0.001

### Folate pathway proteins

Relative concentrations of proteins involved in folate metabolism were measured in brain homogenate collected from adult offspring using Western blots (Fig. 9A-D). SHMT1 immunoreactivity differed between treatment groups (ANOVA, main effect of group, F(2,34)=4.9, η^2^=0.22, p=0.014), with both DPE voles (t-test, t(22)=2.7, d=1.1, p=0.013) and DPE+F voles (t-test, t(24)=3.2, d=1.3, p=0.004) showing elevated SHMT1 relative to controls (Fig. 9A). MTHFR immunoreactivity also differed between groups (ANOVA, main effect of group, F(2,34)=6.0, η^2^=0.26, p=0.006), but with elevated MTHFR in DPE+F voles relative to controls (t-test, Control x DPE+F, t(31)=3.5, d=31, p=0.001) (Fig. 9B). Similarly, FOLR1 immunoreactivity was elevated (ANOVA, main effect of group, F(2,34)=5.3, η^2^=0.24, p=0.010) in DPE+F voles relative to controls (t-test, t(24)=3.1, d=1.2, p=0.005) and DPE voles (t-test, t(22)=2.1, d=0.88, p=0.043) (Fig. 9C). No effects of sex were detected for SHMT1, FOLR1, and MTHFR; and no effects of group or sex were detected for DHFR, MTHFD1, or MTRR (ANOVAs, p>0.05).

**Figure 9.**
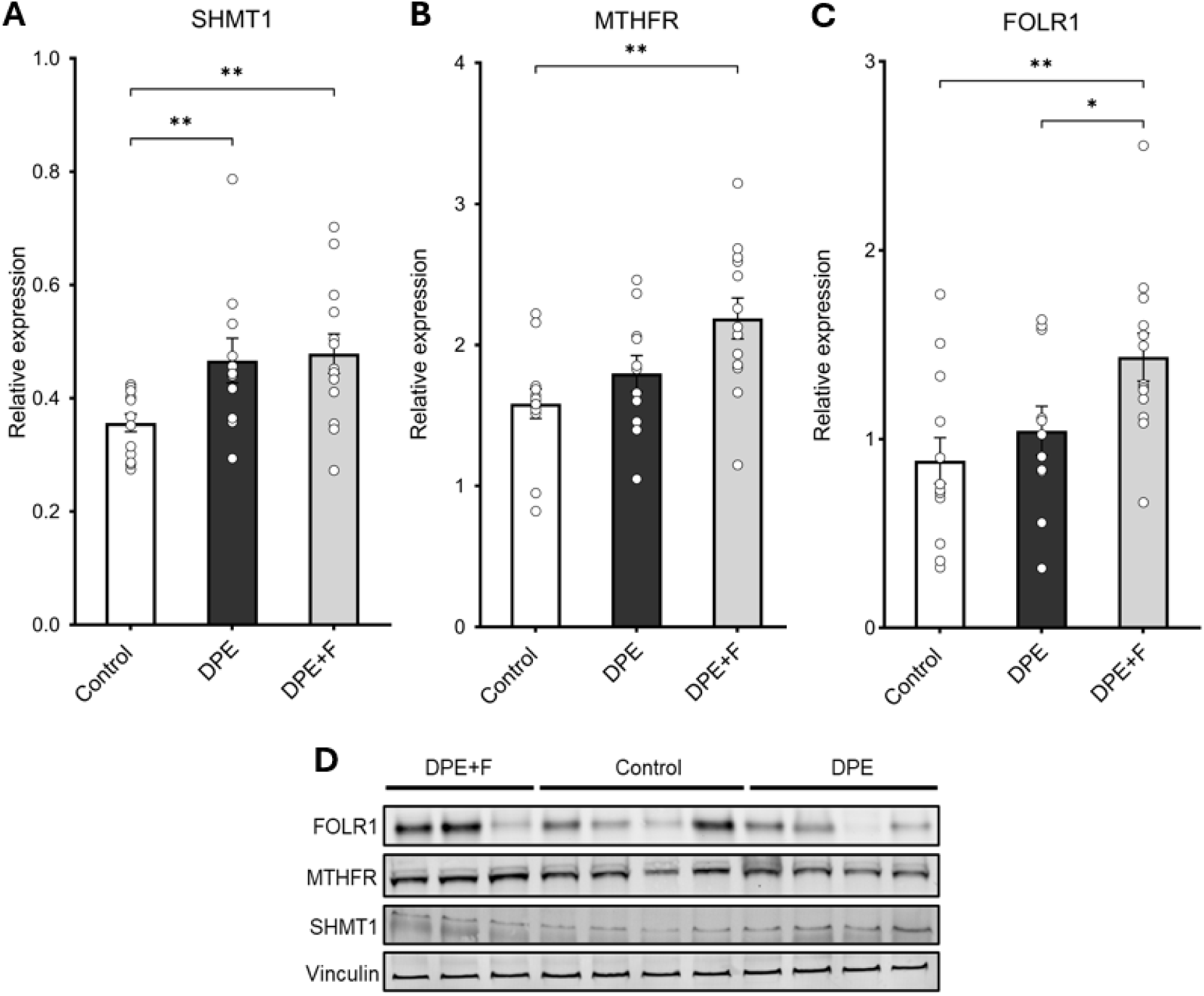
Protein expression of folate metabolism pathway enzymes as measured by immunoreactivity in a Western blot. (A) DPE increased the expression of SHMT1 in DPE voles (N=11) relative to controls (N=13), an effect which was not prevented by maternal folate supplementation (N=13). (B) FOLR1 expression remained unchanged in DPE voles (N=11) compared to controls (N=13) but increased in folate supplemented animals (N=13). (C) MTHFR expression remained unchanged in DPE voles (N=11) compared to controls (N=13) but increased in folate supplemented animals (N=13). (D) Representative bands from Western blots for SHMT1, FOLR1, MTHFR, and the loading control protein vinculin. Error bars are standard error. * p < 0.05 ** p < 0.01 *** p < 0.005 **** p < 0.001

### Individual variability

Prairie voles show significant individual variability at both the behavioral and molecular level, and this variability can be used to find biomarkers that predict either normal variability in behavior or the response to experimental manipulations (Fig. 10A-E, Supplemental Data File 1B) [44]. Across all subjects in the study, individual SHMT1 expression Z-scores were predictive of individual repetitive behavior domain Z-scores, with animals with the highest SHMT1 expression showing the most repetitive behavior (Pearson: R^2^=0.18, p=0.003) (Fig. 10A). This relationship was most evident among DPE voles (R^2^=0.12), suggesting that high SHMT1 expression predicted susceptibility to DPE. In contrast, MTHFR expression was highly predictive of cognitive domain Z-scores across all animals, with the highest MTHFR expression levels corresponding to the best performance on an operant task (Pearson: R^2^=0.21, p=0.031) (Fig. 10B). Plasma folate concentrations were negatively correlated with the cognitive domain (Pearson: R^2^=0.27, p=0.022), with the highest plasma folate corresponding to the worst cognitive scores. Finally, expression of both MTHFR (Pearson: R^2^=0.092, p=0.038) and FOLR1 (Pearson: R^2^=0.13, p=0.012) were positively correlated with repetitive behavior domain Z-scores across all animals, with animals with the highest MTHFR and FOLR1 having the most repetitive behaviors (Figs. 10D-E). This pattern suggests that the individual-level increases in SHMT1 in MTHFR may represent a mechanism of adverse and compensatory group effects, respectively; and that, while increases MTHFR and FOLR1 may have been beneficial relative to behavioral deficits in social behavior, communication, locomotion, and cognition, these increases may have had an adverse effect on individual repetitive behaviors.

**Figure 10.**
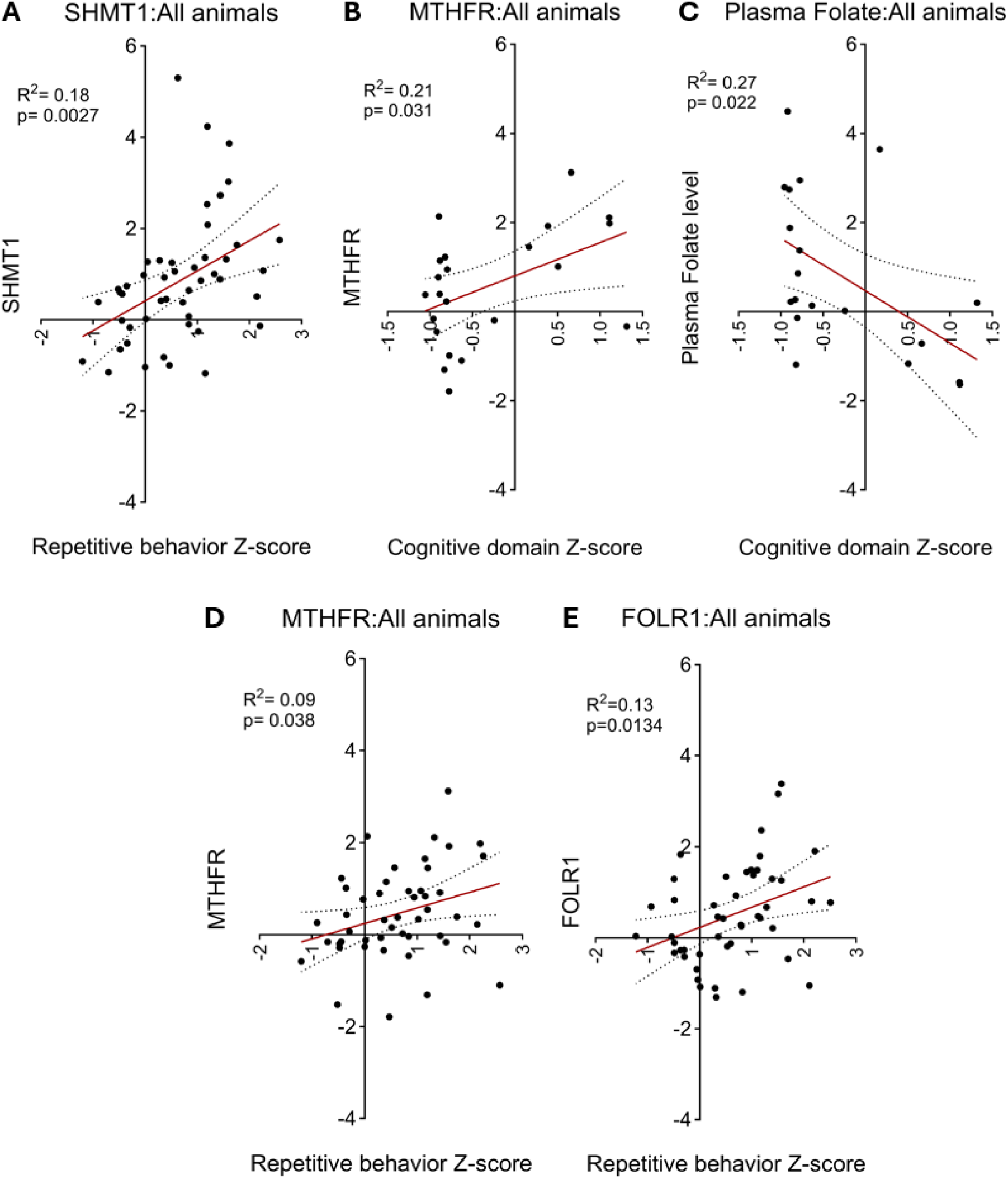
Individual variability in molecular measures predicted individual variability in behavior as represented by domain Z-scores. (A) SHMT1 expression Z-score was positively correlated with the repetitive behavior domain across all experimental animals with both measures (N=47). (B) MTHFR expression Z-score was positively correlated with the cognitive domain across all experimental animals with both measures (N=22). (C) Plasma folate concentration Z-score was negatively correlated with the cognitive domain across all animals with both measures (N=22). Protein expression Z-scores of both (D) MTHFR and (E) FOLR1 were positively correlated with the repetitive behavior domain across all experimental animals with both measures (N=47). The red line represents the line of best fit; curved dotted lines show the 95% confidence intervals.

## DISCUSSION

These experiments were designed to answer two critical questions: whether low-dose developmental pyrethroid exposure in pregnant prairie voles directly affects specific biomarkers of folate metabolism in offspring, and whether high-dose maternal methylfolate supplementation is beneficial in preventing NDD-like behavioral and molecular outcomes. Low-dose DPE in prairie voles caused broad deficits in all five behavioral domains relevant to NDDs. We observed for the first time that DPE disrupts complex social behavior in the prairie vole, specifically the consoling response; whereas DPE was not previously observed to disrupt simple social behaviors in mouse [39]. We also observed that DPE in prairie voles disrupted 24-hour movement patterns, a finding with direct relevance to the disrupted circadian rhythms and sleep disorders that are very highly comorbid with NDDs [62–64]. Most of the deficits induced by DPE were equally present in males and females, although hyperactivity and repetitive rearing were female biased. In combination with human studies implicating DPE as a risk factor for NDDs, and similar behavioral findings in mice [30, 39], this cross-species behavioral evidence strongly reinforces the conclusion that low-dose developmental pyrethroid exposure is a direct cause of NDDs. Low-dose DPE in prairie voles also directly altered folate metabolism in blood and brain, including altering plasma folate concentration and the neural expression of SHMT1 enzyme. Considering the importance of folate in neurodevelopment, this interference with folate metabolism may represent a primary mechanism connecting DPE to adverse developmental outcomes. Most importantly, maternal supplementation with methylfolate (the biologically active form of folate) concurrent with exposures dramatically reduced the overall behavioral phenotype of DPE, specifically by preventing deficits in the social, communication, cognitive, and locomotor domains, but not the deficits in the repetitive behavior domain. These beneficial effects of maternal methylfolate supplementation on the offspring may have occurred through compensatory changes in neural MTHFR and FOLR1 expression and the normalization of plasma and cerebral folate levels. Overall, this evidence supports our hypotheses that DPE in prairie voles interferes with specific aspects of folate metabolism, and that maternal folate supplementation may be an effective strategy in reducing the effects of DPE.

Changes in folate metabolism may represent both an adverse outcome pathway for developmental pyrethroid exposure, and a target for treatment and prevention. It is well known that inadequate folate intake increases the risk for neural tube defects and other developmental disorders, and folate supplementation during pregnancy is an extremely effective preventative treatment [65]. Our findings show that developmental pyrethroid exposure directly alters aspects of folate metabolism, including increased plasma folate concentrations and an increase in SHMT1 expression. Across all animals in our study, individual neural SHMT1 expression was positively correlated with repetitive behavior domain Z-score, with the highest SHMT1 expression corresponding to the most repetitive behaviors; suggesting that increased SHMT1 in DPE voles may represent a toxic response to exposure that drives adverse outcomes. SHMT1 is a critical enzyme in the one-carbon pathway that metabolizes tetrahydrofolate into methylene-tetrahydrofolate (Fig. 1B). SHMT1 also acts as a gate that controls whether one-carbon units (in the form of methylene-tetrahydrofolate) are passed on to the methionine cycle for synthesis of S-adenosyl-methionine (SAM) or are shunted toward nucleotide synthesis to promote cellular growth (Fig. 1B) [66, 67]. The increase in neural SHMT1 expression in DPE voles may cause an imbalance in this gating mechanism, sequestering one-carbon units for use in growth pathways and depleting the supply of one-carbon units for SAM synthesis and subsequent DNA methylation/xenobiotic detoxification reactions [66]. This may partially explain why DPE promotes multimodal changes in molecular pathways for growth in mouse brain transcriptome and proteome [68]. This may be directly relevant to research showing elevated homocysteine, reduced SAM and methionine, and overall reduced methylation capacity in autism [69–71], all of which would be anticipated effects of diversion of one-carbon units away from the methionine cycle. These results justify the critical need for controlled mechanistic experiments evaluating the causal relationship between SHMT1 expression and adverse outcomes in our model.

While maternal methylfolate supplementation did not normalize neural SHMT1 expression, it did drive two potentially compensatory changes. DPE+F voles showed an increase in neural MTHFR expression, and across all animals in this study, individual neural MTHFR expression was positively correlated with cognitive domain Z-score, with the highest MTHFR expression corresponding to the steepest learning curve slopes. This correlation may indicate that MTHFR expression is predicting improved cognition and resilience to DPE. MTHFR, a strong candidate autism risk gene [72], is the enzyme directly downstream of SHMT1 in the one-carbon cycle that converts methylene-tetrahydrofolate into methylfolate. The increase in MTHFR may allow this enzyme to compete with nucleotide synthesis pathways for one-carbon units, potentially compensating for the increase in SHMT1 expression. DPE+F voles also showed an increase in neural FOLR1 expression. As the primary folate transporter in neural cells, the increase in neural FOLR1 may have compensated for SHMT1-driven folate sequestration by allowing cells to import more of the direct precursor for the methionine cycle and feeding the entire one-carbon cycle more total one-carbon units. This increased cellular import of folate could also explain the normalization of plasma folate concentrations in DPE+F voles; and our whole-brain samples included choroid plexus, where FOLR1 is highly expressed and transports folate directly from plasma into the brain. Nonetheless, individual MTHFR expression levels were positively correlated with repetitive behavior domain Z-scores. Therefore, while compensatory increases in MTHFR may have driven the positive benefits of maternal methylfolate supplementation in the social, communication, cognitive, and locomotor domains, these compensatory changes may also have contributed to the increase in repetitive behaviors in DPE+F voles. The potential of these mechanisms in exposed animals to both compensate for DPE, and to drive increases in repetitive behaviors, needs to be tested with controlled experiments.

In adulthood, DPE voles showed higher plasma folate concentrations, which were normalized by maternal methylfolate supplementation. The CLIA assay does not differentiate between forms of folate, making this an aggregate measure of all folates present. Since high plasma folate concentrations in the voles predicted low cognitive domain scores, this leads to the perplexing conclusion that higher concentrations of folate in the plasma may be adverse. Differences in dietary consumption are not likely to be the source of these group differences in plasma folate, since our laboratory prairie voles all consume the same commercial diet, containing 7.4 ppm folate that is 81% in synthetic folic acid form [51]. Based on our estimates, the average prairie vole on this diet is consuming 0.5 mg/kg folate per day -- about 60 times the amount recommended for humans – the vast majority of which is in folic acid form. Increased plasma folate in DPE voles may instead be a biomarker of poor folate metabolism, attributable to either incomplete conversion to methylfolate in the liver [73, 74] (leading to accumulation of unmetabolized folic acid in plasma), or to malabsorption of folate into tissues. This could help to explain why methylfolate supplementation was still beneficial despite the already large dietary intake of folic acid in prairie voles.

Due to successes in promoting folate supplement use and the fortification of foods in the United States, unmetabolized folic acid is present in 95% of serum samples from the general population [75]. Studies addressing whether unmetabolized folic acid itself is harmful have been inconsistent [75–81], and the current scientific consensus is that unmetabolized folic acid levels in humans also represents a biomarker of poor folate metabolism, which could pose health risks and increase the incidence of NDDs [76]. The developmental exposure model in prairie voles, as a model of disrupted folate metabolism relevant to NDDs, is ideally positioned to address this important and poorly understood scientific issue in future studies.

Among the wide-ranging effects of developmental pyrethroid exposure we observed, the impact on the cognitive domain is perhaps the most concerning. In humans, pyrethroid exposure during development has been identified as a risk factor for developmental delay, learning disabilities, and reduced intelligence [22, 30, 33, 35, 82–85] and similar effects have even been observed in chronically exposed adults [86]. In our study, prairie voles exposed developmentally to a low dose of deltamethrin failed to learn the most basic operant task available, nose poking for food when hungry. While developmentally exposed prairie voles performed normally on a fear conditioning task, mice with the same developmental exposure failed to associate a tone with a threat, one of the most basic fear learning tasks available [39]. This current research adds a third species to the extensive literature in rats and mice showing cognitive deficits caused by pyrethroid exposures during development [30, 39, 87–90] and across the lifespan [91–94]. While exposure doses in these animal studies vary greatly, the developmental exposure doses required to produce measurable reductions in cognitive performance were quite low, on par with the dose used in this study. The totality of evidence strongly supports the conclusion that low-dose developmental pyrethroid exposure causes reduced cognitive function in humans and animals.

Our study incorporated several key strengths intended to increase rigor and reproducibility in research relevant to neurodevelopmental disorders [2]. First and foremost, we chose the prairie vole as our animal model. The prairie vole has been extensively used in behavioral neuroscience to study brain mechanisms of social behavior, owing to their species-specific complex social behaviors; and their outbred genetics provides an opportunity to discover individual biological variability that predicts or drives individual behavioral variability [44]. Our intuition that complex social behaviors in the prairie vole would be more sensitive to disruption than the relatively simple social behaviors of mice appears to have been justified. We used a broad behavioral domain approach, testing for behaviors relevant to five domains affected in neurodevelopmental disorders [50]. Multiple outcomes were used for most domains to decrease task-specific bias, including outcomes across the lifespan; and many of the specific assays were selected to replicate findings from the mouse [30, 39]. We used maximally automated scoring methods, a mix of both sexes, and behaviorally naïve animals for molecular assays, all to reduce bias and eliminate sources of systematic variability. Finally, we used the litter-based design: because the primary exposure is in the dam, the litter is considered the experimental unit, and each subject derives from a unique litter.

There are several important limitations that should be considered when interpreting our results. One major limitation of our study is the use of a single exposure dose of deltamethrin. The developmental dose of 3 mg/kg was chosen based on our previous studies as one sufficient to alter NDD-relevant mouse behavior and alter folate metabolism in mouse brain [39, 42, 68]. Nonetheless, prior literature recognizes this dose as the maternal toxicity “no observable adverse effect level” (NOAEL), and it is well below both the developmental NOAEL of 12 mg/kg, and the regulatory benchmark dose (14.5 mg/kg) set by the EPA [43, 95]. These prior determinations highlight the fact that the developmental exposure dose we used in this study was previously assumed to produce few or no effects. In our study, we did not measure levels of deltamethrin in offspring because their last exposure was at weaning age. To put the maternal exposure levels in context, studies in pregnant mice exposed to 1.5 mg/kg of permethrin had serum concentrations of 260 ng/mL [191]. In another study, pregnant Chinese women had an average of 39 ng/mL deltamethrin in their blood [192], while the level of deltamethrin in the general population of Beijing ranged from 0-17.34 ng/ml [193]. In the U.S., urinary metabolite levels rose from 0.292 ng/mL [194] in 1999–2000 to 660 ng/mL in 2011–2012 [195]. These statistics suggest that the chosen exposure dose in this study may reflect an above average exposure yet remains below the established developmental no observable adverse effect level and the EPA’s benchmark dose.

Other limitations include the use of a single supplemental dose of methylfolate and the absence of a “methylfolate only” group. This experimental design limits our ability to draw conclusions about the appropriate dose of supplementation or the appropriate form (i.e., methylfolate vs. folic acid). Nonetheless, the fact that voles in all three experimental groups were consuming high levels of folic acid in their laboratory diet suggests that supplementation with a biologically active form may have been critical to the observed effects. Importantly, in our study, methylfolate supplementation was only ever given in combination with deltamethrin exposure, and therefore the effects of methylfolate supplementation can only be understood as the combined effect of exposure and supplementation. Thus, we are unable to conclude from our data whether the increase in repetitive behaviors in DPE+F voles were a combined effect of exposure and supplementation or were instead a primary effect of high-level methylfolate supplementation. Similarly, we cannot conclude whether the positive effects of methylfolate in DPE+F voles represent specific compensation for the effects of DPE or non-specific benefits of supplementation. In terms of proteins, we focused on the primary folate transporter in brain and selected major enzymes of the one-carbon cycle, as opposed to using a broader proteomics approach that could have detected a wider range of changes relevant to folate metabolism. We also used a whole-brain approach, which cannot inform on any specific brain regions impacted by DPE. All molecular samples were collected from adult offspring, so we are unable to conclude at what point during development the molecular changes occurred; although we do see the earliest behavioral deficits emerge as early as 1 week of age. Finally, the elevated plasma folate levels seen in DPE voles are unlikely to have been caused by the observed changes in folate metabolism in brain, and instead point to an as-of-yet unexplored peripheral effect of DPE on folate metabolism or absorption, possibly in the liver.

## CONCLUSIONS

Our study provides causal evidence in strong support of the conclusion that low-dose chronic pyrethroid exposure during pregnancy is a direct cause of neurodevelopmental disorders. Developmental pyrethroid exposure directly alters folate metabolism, which may provide a mechanism for some or all of the developmental effects on behavior. This study provides causal evidence (consistent with human epidemiology [41]) supporting the conclusion that high-dose maternal folate supplementation before and during pregnancy helps to compensate for the detrimental effects of pyrethroid exposure; however, current guidelines may be suggesting both the wrong dose, and the wrong type, of folate supplementation. Based on the totality of evidence, we highly recommend daily consumption of 800 µg of folate both prior to and during pregnancy, to help protect against exposure to pyrethroids; and we specifically recommend that, in addition to the currently recommended 400 µg of daily folic acid from prenatal supplements, women take an additional 400 µg in a natural bioactive form, such as methylfolate supplements and/or foods naturally high in folate. Finally, it is to be noted that methylfolate does not appear to counteract developmental pyrethroid exposure, but only partially compensates for it, thereby emphasizing the importance of reducing exposure to pyrethroids during pregnancy.

## Supporting information

Suppemental Data File 1

## DATA AVAILABILITY

All raw data and metadata are provided as supplemental materials.

## SUPPLEMENTAL MATERIALS

Supplemental figures and tables can be found in the Supplemental Materials file. Raw data in the form of spreadsheets can be found in Supplemental Data File 1.

## ACKNOWLEDGMENTS

We acknowledge the Cornell Vole Core as the source of our prairie vole breeding stock, which is itself sourced from wild prairie voles trapped at Illinois field sites by Dr. Alex Ophir. Prairie vole research as it exists today could not continue without Dr. Ophir’s ongoing heroic efforts.

## AUTHOR CONTRIBUTIONS

NS and JPB designed the overall research study, including the conceptual background and experimental strategy. FSH designed the operant conditioning experiments. Behavioral and molecular experiments were performed by, and data collected and analyzed by, NS, IH, SP, AEN, RL, SPK, CNN, IM, DH, JAC, and KKZ, under the direction and supervision of NS and JPB. Operant conditioning experiments were performed by AT, SEB, SPD, and EMC under the direction and supervision of FSH. RS, SG, and JV performed additional data analysis. ZVJ provided additional analysis and interpretation of partner preference data. MLG provided additional analysis and interpretation of vocalization data. KLN provided experimental, administrative, and managerial support. NS prepared the final figures. NS and JPB interpreted all results and prepared the initial draft of the manuscript. All authors participated in writing the manuscript and approved the final version.

