## Supplemental Materials for "Folate prevents the autism-related phenotype caused by developmental pyrethroid exposure in prairie voles"

**This PDF file includes:**

Supplemental Tables S1-S2

Supplemental Figures S1

| **Experimental use** | **Control** | **DPE** | **DPE+F** |
| --- | --- | --- | --- |
| **Total** | N=18 litters | N=17 litters | N=16 litters |
| **Behavior:** |  |  |  |
| Ultrasonic vocalizations | N=18 | N=17 | N=16 |
| Observation | N=17 | N=16 | N=14 |
| Marble burying | N=17 | N=16 | N=14 |
| Locomotion | N=17 | N=16 | N=14 |
| 24-hour mobility | N=17 | N=16 | N=14 |
| Partner preference | N=12 | N=12 | N=12 |
| Consoling behavior | N=12 | N=12 | N=12 |
| Fear conditioning | N=12 | N=12 | N=12 |
| Operant conditioning | N=12 (+cohort 2, N=10) * | N=14 (+cohort 2, N=13) * | N=14 |
| **Molecular:** |  |  |  |
| Chemiluminescence | N=14 | N=14 | N=14 |
| Western blot | N=18 male, N=18 female | N=17 male, N=17 female | N=16 male, N=16 female |

**Table S1.** Allocation of subject offspring to experimental tests prior to task-specific eliminations.

* Subjects included from a separate, independent cohort.

| **NDD-relevant domain** | **Domain-relevant behavior assay** | **Deficit observed in DPE mouse [30, 39]** | **Primary outcomes** | **Descriptive outcomes** |
| --- | --- | --- | --- | --- |
| Social domain | Partner preference test  Consoling behavior test | **Not tested**  **Not tested** | Huddling time  Partner grooming * |  |
| Communication domain | Ultrasonic vocalizations | Yes | Call count * | Call duration  Principal frequency  Peak frequency  Delta frequency  Slope  Sinuosity  Tonality |
| Cognitive domain | Operant conditioning  Fear conditioning | Yes  Yes | Task acquisition  Freezing (recall) | 10-day learning curve  Learning curve slope *  Freezing (acquisition)  Freezing (extinction) |
| Locomotor domain | Locomotor activity  24-hour mobility | Yes  **Not tested** | Total locomotion *  Locomotion over time | Locomotion (ZG7-8) * |
| Repetitive behavior domain | Marble burying  Self-grooming  Rearing | Yes  Yes  Yes | Marbles buried *  Grooming duration *  Rearing duration * |  |

**Table S2.** Assays selected to represent behavioral domains relevant to neurodevelopmental disorders.

* Outcome measures selected for inclusion in the Combined Phenotype Z-score calculation.

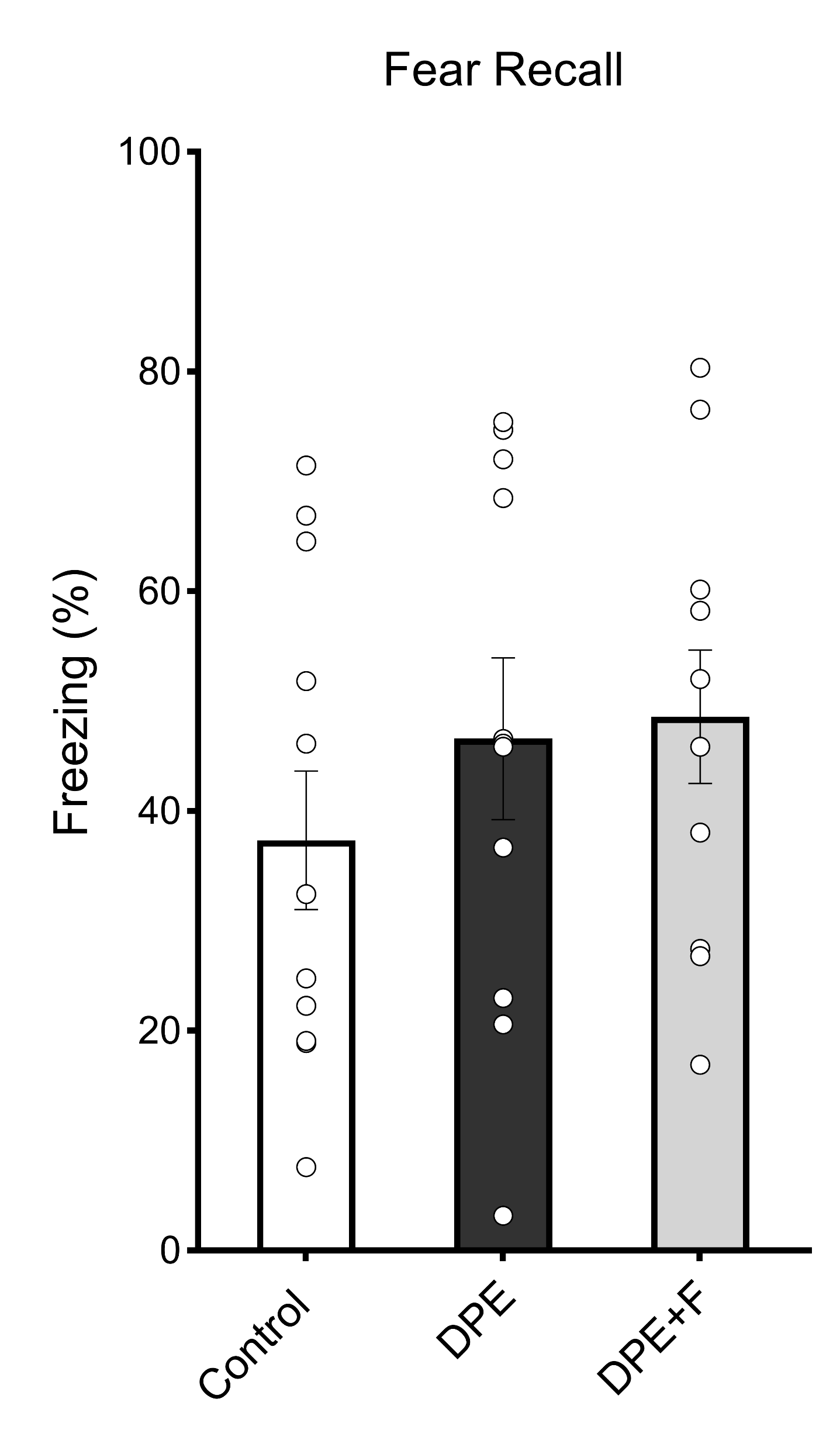

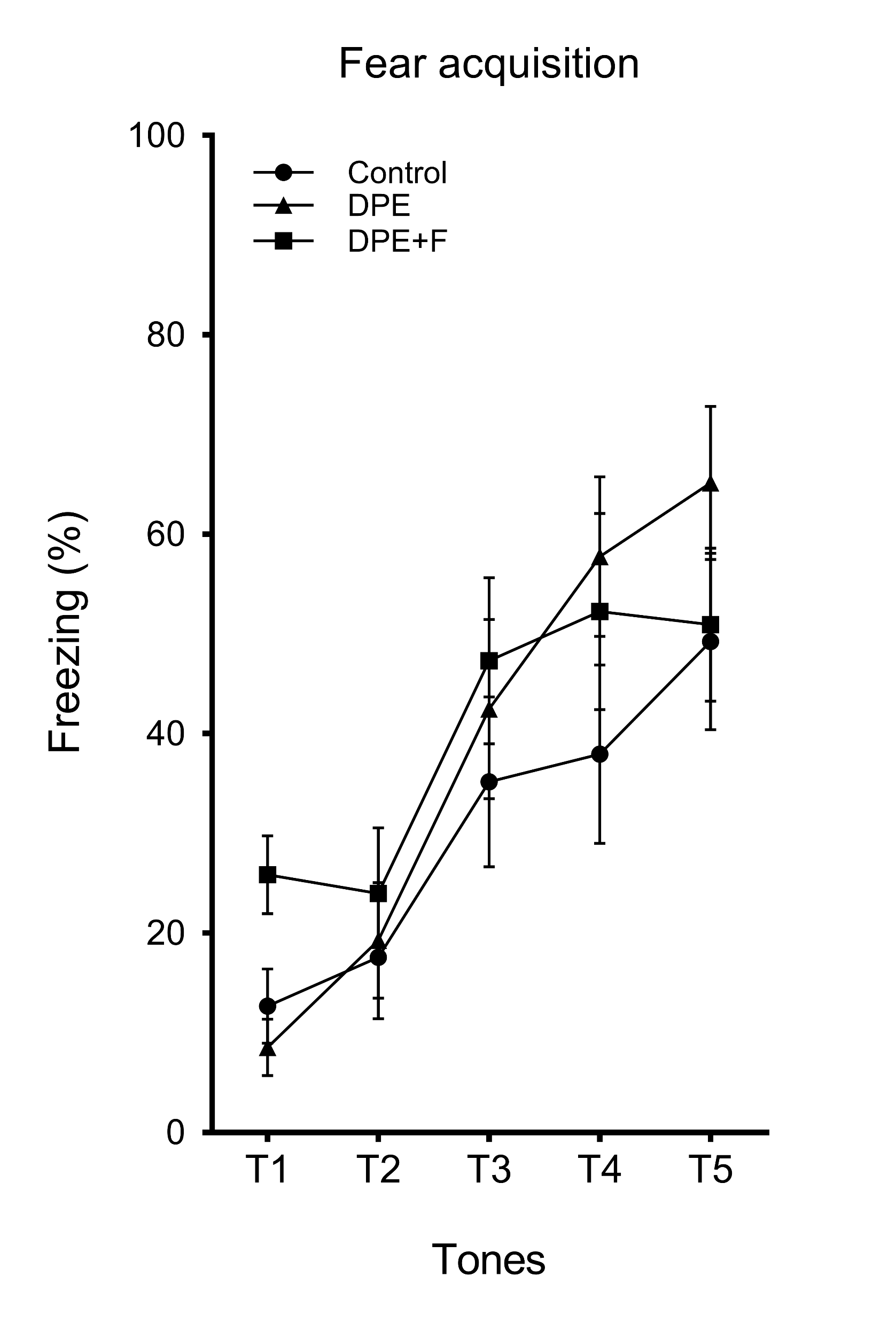

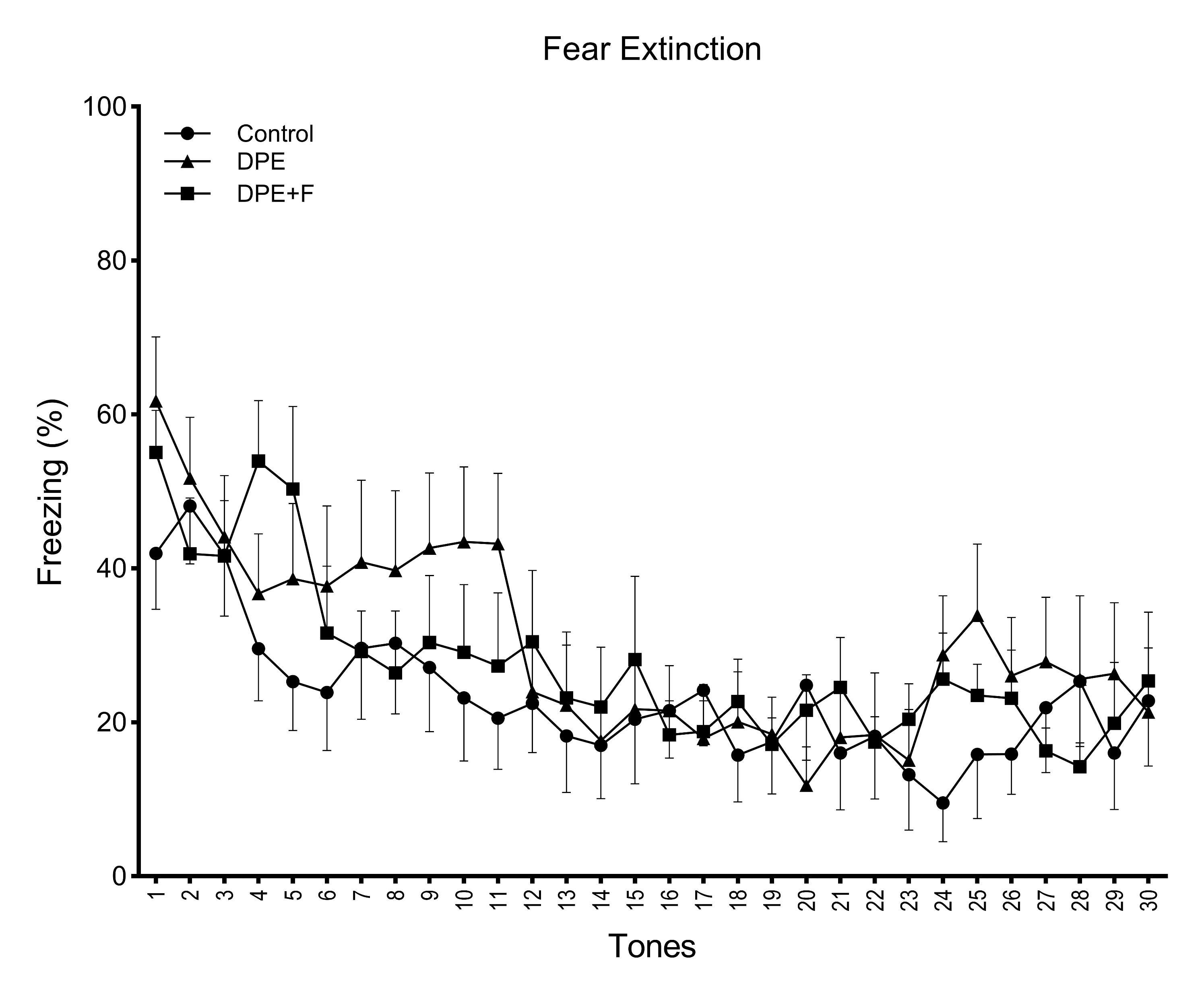

**A**

**B**

**C**

**Figure S1.** Fear conditioning in prairie voles. A) Acquisition of the freezing response during classical fear conditioning with tone-shock pairings. B) Recall of the tone-associated freezing response in the absence of shocks on the second day. C) Extinction of the tone-associated freezing response across 30 tones on days 2 and 3.
